# *rpoB*, a promising marker for analyzing the diversity of bacterial communities by amplicon sequencing

**DOI:** 10.1101/626119

**Authors:** Jean-Claude Ogier, Sylvie Pagès, Maxime Galan, Mathieu Barret, Sophie Gaudriault

**Affiliations:** INRA, Université de Montpellier, UMR1333-DGIMI, 34095 Montpellier Cedex 05, France; INRA, AGROCAMPUS-Ouest, Université d’Angers, Beaucouzé, France; CBGP, INRA, CIRAD, IRD, Montpellier SupAgro, Univ. Montpellier, 755 avenue du Campus Agropolis, CS 300 16, F-34988 Montferrier sur Lez cedex, France

**Author notes:** Corresponding author: Jean-Claude OGIER, UMR 1333 DGIMI, Faculté des Sciences, Place Eugène Bataillon, CC 54, 34095 MONTPELLIER CEDEX 05, FRANCE, 77. Email addresses of the authors.

**Keywords:** *rpoB*, 16S rRNA gene, metabarcoding, mock communities, entomopathogenic nematodes

## Abstract

**Background:** Microbiome composition is frequently studied by the amplification and high-throughput sequencing of specific molecular markers (metabarcoding). Various hypervariable regions of the 16S rRNA gene are classically used to estimate bacterial diversity, but other universal bacterial markers with a finer taxonomic resolution could be employed. We compared specificity and sensitivity between a portion of the *rpoB* gene and the V3V4 hypervariable region of the 16S rRNA gene.

**Results:** We first designed universal primers for *rpoB* suitable for use with Illumina sequencing-based technology and constructed a reference *rpoB* database of 45,000 sequences. The *rpoB* and V3V4 markers were amplified and sequenced from (*i*) a mock community of 19 bacterial strains from both Gram-negative and Gram-positive lineages; (*ii*) bacterial assemblages associated with entomopathogenic nematodes. In metabarcoding analyses of mock communities with two analytical pipelines (FROGS and DADA2), the estimated diversity captured with the *rpoB* marker resembled the expected composition of these mock communities more closely than that captured with V3V4. The *rpoB* marker had a higher level of taxonomic affiliation, a higher sensitivity (detection of all the species present in the mock communities), and a higher specificity (low rates of spurious OTU detection) than V3V4. We applied both primers to infective juveniles of the nematode *Steinernema glaseri*. Both markers showed the bacterial community associated with this nematode to be of low diversity (< 50 OTUs), but only *rpoB* reliably detected the symbiotic bacterium *Xenorhabdus poinarii*.

**Conclusions:** Our results confirm that different microbiota composition data may be obtained with different markers. We found that *rpoB* was a highly appropriate marker for assessing the taxonomic structure of mock communities and the nematode microbiota. Further studies on other ecosystems should be considered to evaluate the universal usefulness of the *rpoB* marker. Our data highlight two crucial elements that should be taken into account to ensure more reliable and accurate descriptions of microbial diversity in high-throughput amplicon sequencing analyses: i) the need to include mock communities as controls; ii) the advantages of using a multigenic approach including at least one housekeeping gene (*rpoB* is a good candidate) and one variable region of the 16S rRNA gene.

## Background

The recent emergence of high-throughput sequencing platforms has revolutionized the study of complex microbial communities. Many of these studies involve the PCR amplification and sequencing of a taxonomic marker from complex communities of organisms. The sequences obtained can then be compared with databases of known sequences to identify the taxa present in the microbial community. The 16S rRNA gene is the most common marker used for this purpose in bacterial ecology, particularly as exhaustive reference databases have been compiled for this taxonomic marker: the Greengenes database [1], the Ribosomal Database Project (RDP) [2], and SILVA [3], for example. In addition to their extensive catalogs of curated 16S rRNA gene sequences, each of those portals also offers a series of tools for sequence analysis. However, 16S rRNA markers are also known to be the major source of bias in the amplicon sequencing approach [4]. Estimates of microbial diversity are generally biased by the variable number and sequence heterogeneities of 16S rRNA operons in bacterial species, generally leading to an overestimation of species richness [5, 6]. Furthermore, a growing number of publications have reported limitations to the use of the 16S rRNA gene in ecological studies due to its poor discriminatory power in certain bacterial genera [7], resulting in poor discrimination between species. In light of these major drawbacks, alternative, or at least complementary taxonomic markers should be sought for metabarcoding projects.

A few studies have tested other taxonomic markers by targeting conserved protein-coding genes, also known as housekeeping genes. Like the 16S rRNA gene, housekeeping genes are essential and ubiquitous genes universally present in the bacterial kingdom. However, these housekeeping genes evolve much more rapidly than the16S rRNA gene, and are therefore useful for differentiating between lineages that have recently diverged [8]. Moreover, these housekeeping genes are generally present as single copies in bacterial genomes, limiting the overestimation of operational taxonomical units (OTUs) in microbial assemblages. Despite these advantages, they are very rarely used in metabarcoding analyses, mostly due to the lack of an exhaustive sequence database for these genes. The protein-coding genes that have been tested for the assessment of microbial diversity include the genes for DNA gyrase subunit B (*gyrB*) ([8-10], RNA polymerase subunit B (*rpoB*) [7, 11, 12], the TU elongation factor (*tuf*) [13], the 60 kDa chaperonin protein (*cpn60*) [14].

The objective of this study was to assess the potential of the *rpoB* gene as an alternative universal phylogenetic marker for metabarcoding analysis. The *rpoB* gene has a number of potential advantages. Previous reports have shown this marker to be suitable for phylogenetic analyses, as it provides a better resolution at species level than the 16S rRNA gene [15-18]. Moreover, the *rpoB* gene is sufficiently long (4026 nc) for the rational identification of conserved genomic regions for the design of universal primers for the amplification of short barcodes (reads < 300 nc). Specific *rpoB* primer sets targeting Proteobacteria have been designed for high-throughput sequencing studies [12], but universal *rpoB* primers have not been tested in next-generation sequencing studies.

We designed universal *rpoB* primers suitable for use with Illumina sequencing-based technology and we constructed an *rpoB* reference database of 45,000 sequences. The specificity and sensitivity of *rpoB* were assessed on a mock bacterial community composed of DNA extracted from 19 strains, and compared with those for the V3V4 regions of the 16S rRNA gene. We also applied this community profiling approach to infective juveniles (IJs) of *Steinernema glaseri*, an entomopathogenic nematode [19, 20]. IJs are specifically associated with the intestinal bacterium *Xenorhabdus poinarii* (Enterobacteriaceae) [21, 22], which is therefore an effective control for evaluating the performance of taxonomic markers for Illumina sequencing analyses. We found that bacterial richness, in both mock communities and the nematode microbiota, was estimated more accurately with the *rpoB* marker than with the 16S rDNA marker, confirming the advantage of *rpoB* as an alternative or complementary marker to the traditional variable regions of the 16S rRNA gene.

## Results

### Comparison of *rpoB* and V3V4 markers distinguishing by Sanger sequencing 19 individual taxa

We compared the taxonomic discrimination potentials of the *rpoB* and V3V4 markers, by extracting genomic DNA from the individual bacterial strains used to constitute the mock communities; then amplifying and Sanger sequencing the *rpoB* gene fragment (∼430 bp) and the V3V4 region of the 16S rRNA gene (∼450 bp). We used the RDP classifier tool (sequence similarity threshold = 97%, bootstrap confidence cutoff = 80%) to assign the sequences to taxa (Table 1). Taxonomic affiliation was determined more precisely with the *rpoB* gene fragment than with the V3V4 region, because 13 and 0 OTUs were affiliated to a species with *rpoB* and V3V4, respectively (Table 1 and Figure 1A, lanes “Expected-*rpoB* or Expected-16S”). Moreover, the *rpoB* marker had a higher sensitivity than the V3V4 marker (19 OTUs *versus* 17 OTUs, Table 1).

**Table 1.**
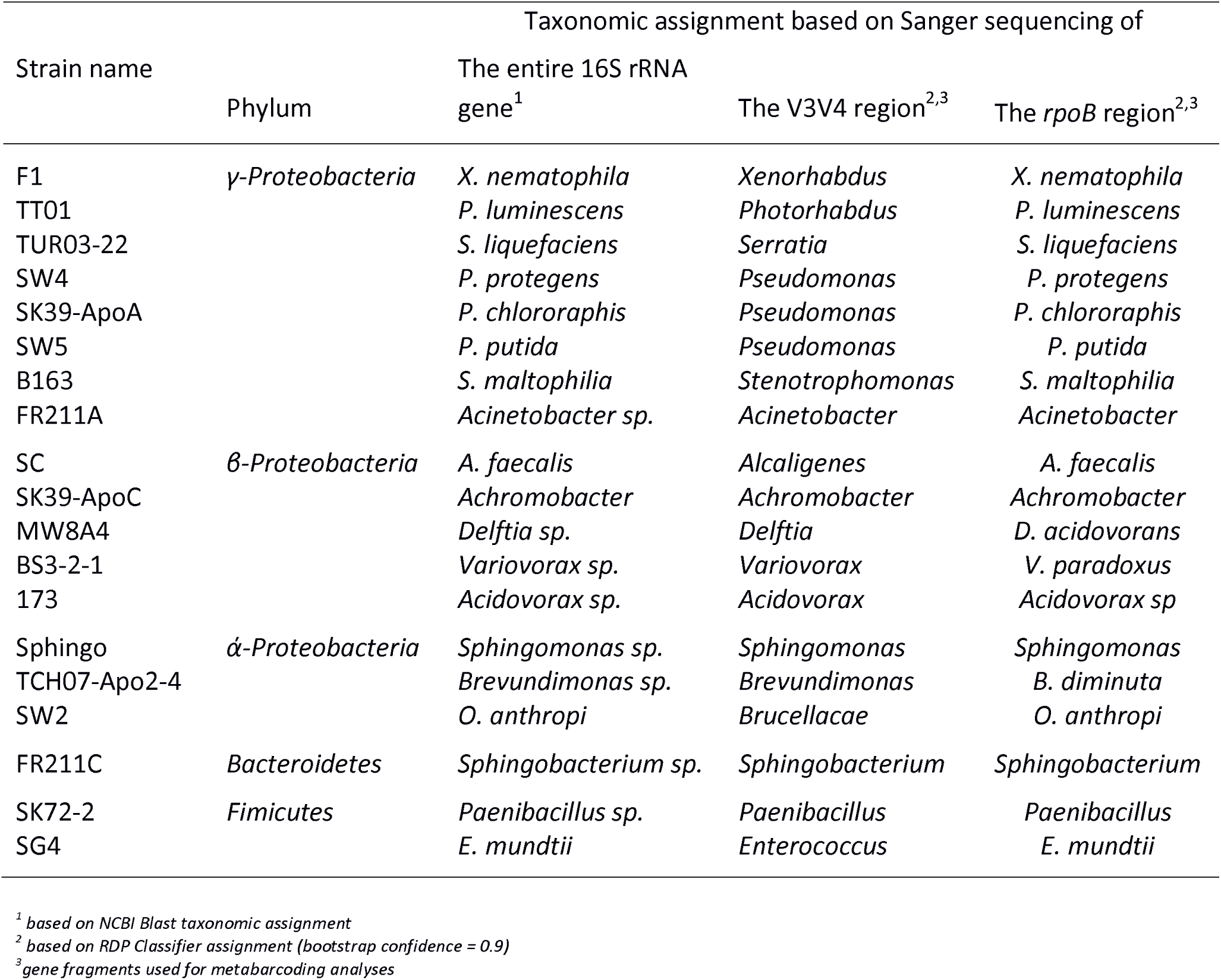
List of strains used for the extraction of the DNAs used to constitute the mock communities

**Figure 1.**
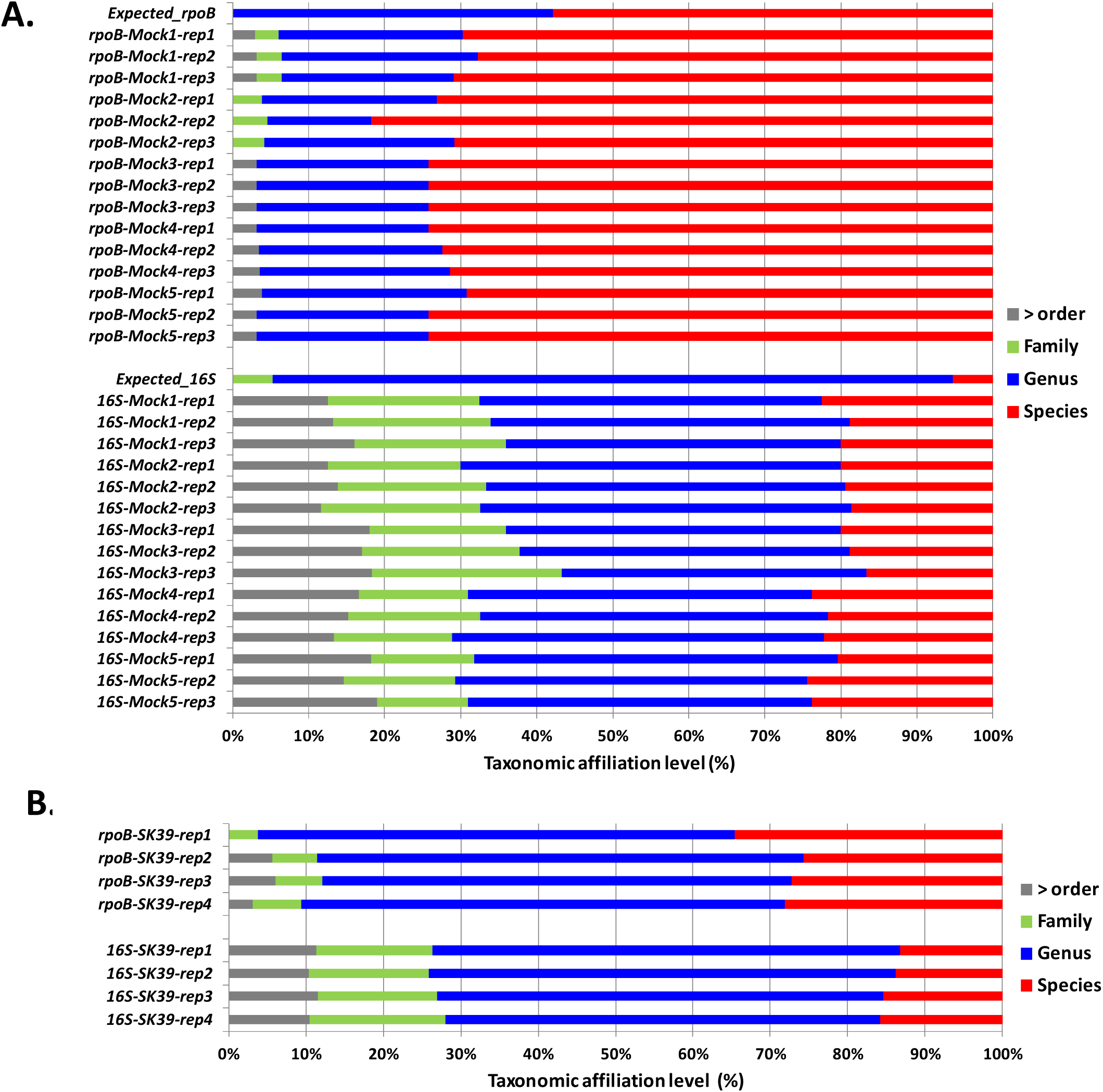
Impact of the metabarcode markers on OTU taxonomic affiliations. Histogram showing the percentage of OTU affiliations at different taxonomic ranks (*x*-axis) for *rpoB* and 16S Illumina-amplicon sequencing for **(A)** the 15 mock community samples (five mock communities and 3 replicates per mock community) **(B)** the microbiota of the nematode *Steinernema glaseri* SK39 (four replicates). Gray, green, blue and red histogram bars correspond to the taxonomic ranks of order (and higher levels), family, genus and species, respectively. The total number of OTUs observed after sequence analysis via the FROGS pipeline was included in the analysis. The *Expected_rpoB* and *Expected_16S* lanes correspond to the taxonomic affiliations determined by Sanger sequencing of each of the taxa making up the mock communities.

### Comparison of the *rpoB* and V3V4 markers for metabarcoding descriptions of the bacterial community making up five artificial mock communities

We constituted five different mock communities differing in the proportions of two taxa, *Xenorhabdus nematophila* and *Photorhabdus luminescens*. The proportions of the 19 bacterial species making up the five mock communities, in terms of genome equivalents, are shown in Table 2. We performed metabarcoding analyses of the five mock communities (three technical replicates per mock community) with both *rpoB* and the V3V4 region of the 16S rRNA gene (referred to hereafter as 16S). The targets were amplified and sequenced, and the sequences were processed with two different pipelines, FROGS (Find Rapidly OTUs with Galaxy Solution) and DADA2, to minimize bias related to the sequence analysis process. For the *rpoB* region, we obtained 224,129 reads (mean of 14,941 reads per sample) and 251,963 reads (mean of 16,797 reads per sample) after FROGS and DADA2 processing, respectively (Additional file 1). For the 16S marker, we obtained 266,494 reads (mean of 17,766 reads per sample) and 206,641 reads (mean of 13,776 reads per sample) after FROGS and DADA2 processing, respectively (Additional file 1). Rarefaction curves obtained with quality-filtered reads indicated that sequencing depth was sufficient for both 16S and *rpoB* (Additional file 2). A comparison of the taxonomic affiliation levels of the observed OTUs (Figure 1A) confirmed that the *rpoB* gene fragment had the higher taxonomic discriminating potential; 60 to 80% of OTU assignations were at the species level for *rpoB* marker, but only 20% for the 16S marker (Figure 1A). We also noted that numerous OTUs (30 to 40%) could not be assigned to a finer taxonomic level than the family with the 16S marker.

**Table 2.** Composition of the five mock communities

| Bacterial species name | Mock1<sup>a</sup> | Mock2<sup>a</sup> | Mock3<sup>a</sup> | Mock4<sup>a</sup> | Mock5<sup>a</sup> |
| --- | --- | --- | --- | --- | --- |
| *Enterococcus mundtii* | 0.09 | 0.01 | 0.17 | 0.17 | 0.17 |
| *Alcaligenes faecalis* | 1.58 | 0.11 | 2.81 | 2.91 | 2.93 |
| *Variovorax paradoxus* | 2.29 | 0.16 | 4.08 | 4.23 | 4.25 |
| *Pseudomonas protegens* | 2.07 | 0.14 | 3.69 | 3.83 | 3.85 |
| *Stenotrophomonas maltophilia* | 8.39 | 0.57 | 14.93 | 15.50 | 15.57 |
| *Brevundimonas diminuta* | 6.15 | 0.42 | 10.94 | 11.36 | 11.41 |
| *Serratia liquefaciens* | 1.45 | 0.10 | 2.58 | 2.68 | 2.69 |
| *Acinetobacter sp.* | 6.21 | 0.42 | 11.05 | 11.47 | 11.52 |
| *Pseudomonas chlororaphis* | 3.10 | 0.21 | 5.51 | 5.72 | 5.74 |
| *Ochrobactrum anthropi* | 2.15 | 0.15 | 3.83 | 3.98 | 4.00 |
| *Pseudomonas putida* | 3.11 | 0.21 | 5.53 | 5.74 | 5.77 |
| *Sphingobacterium sp.* | 5.72 | 0.39 | 10.17 | 10.56 | 10.60 |
| *Acidovorax sp.* | 2.24 | 0.15 | 3.98 | 4.14 | 4.15 |
| *Achromobacter sp.* | 1.42 | 0.10 | 2.53 | 2.63 | 2.64 |
| *Paenibacillus sp.* | 3.21 | 0.22 | 5.72 | 5.93 | 5.96 |
| *Delftia sp.* | 1.41 | 0.10 | 2.51 | 2.61 | 2.62 |
| *Sphingomonas sp.* | 3.28 | 0.22 | 5.83 | 6.05 | 6.08 |
| *Xenorhabdus nematophila* | 35.87 | 36.69 | 3.35 | 0.40 | 0.04 |
| *Photorhabdus luminescens* | 10.24 | 59.64 | 0.80 | 0.08 | 0.01 |

<sup>a</sup> percentage based on genome number equivalents

For each sample mock community replicate, OTU numbers after the application of two thresholds to individual sample read abundances (0.1% and 1%) are indicated in Additional file 1. We evaluated the impact of the use of the *rpoB* and 16S markers on the number of OTUs detected in metabarcoding analyses (Figure 2). The numbers of OTUs detected depended on the marker, and were generally greater for the 16S than for the *rpoB* marker; they also depended on the analysis pipeline. Interestingly, depending on the abundance threshold used, both markers either overestimated (cutoff = 0.1%) or underestimated (cutoff = 1%) the number of OTUs.

**Figure 2.**
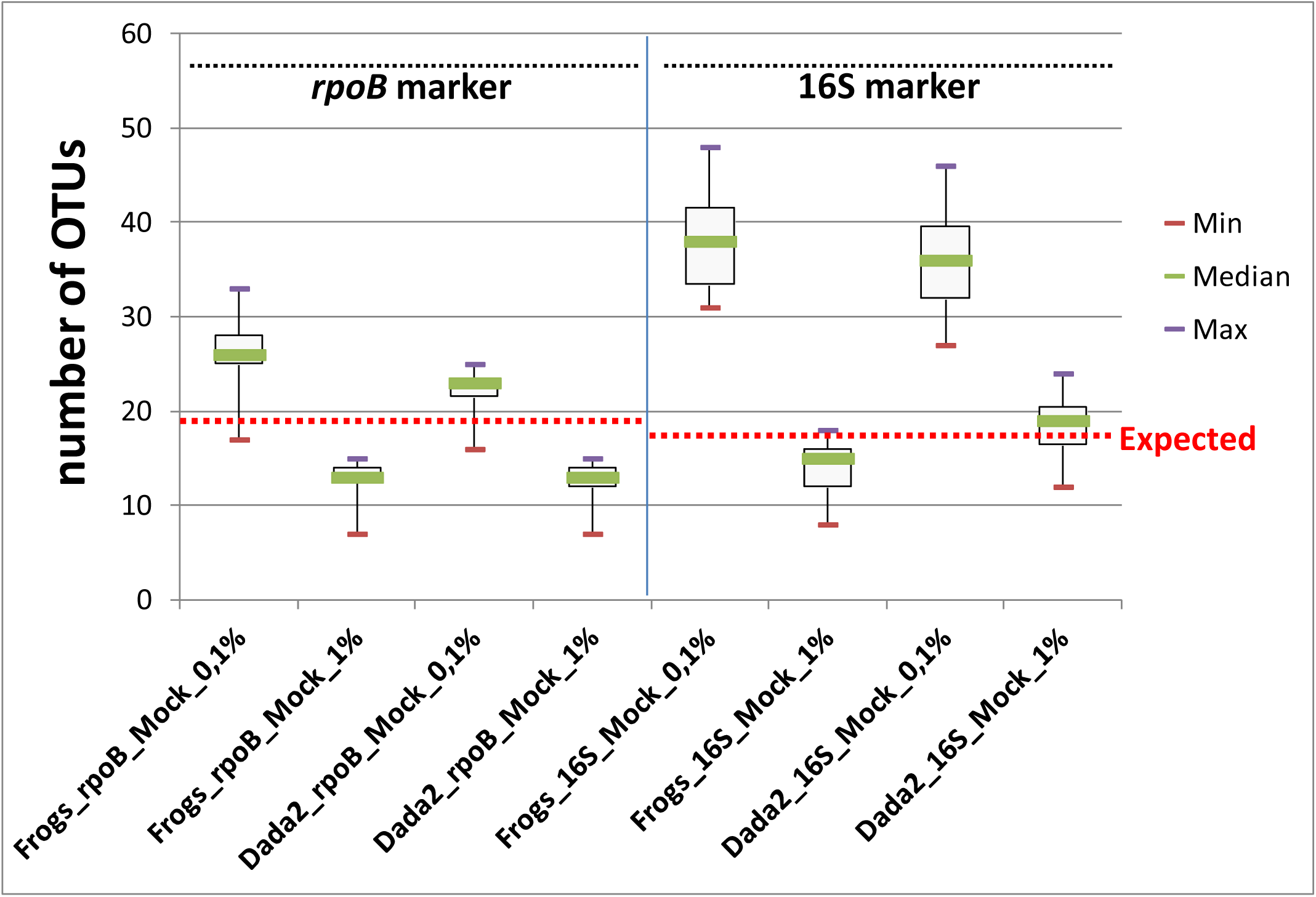
Impact of the markers, sequence analysis pipeline tools and filtering method on observed OTU richness after Illumina-amplicon sequencing of the five mock communities. Boxplots represent the variation of OTU numbers as a function of: i) Illumina-amplicon sequencing procedure, based on the *rpoB* marker or the 16S marker; ii) sequence analyses process, based on the FROGS or DADA2 pipeline; iii) OTU filtering method based on a 0.1% read number threshold or a 1% read number threshold. Each boxplot corresponds to a statistical analysis of the 15 mock community samples (five mock communities, three replicates per mock community); thin purple and red lines correspond to minimum and maximum values, respectively; and the thicker green lines in the boxes correspond to the medians. Red dashed lines correspond to the expected number of OTUs in the mock communities (19 and 17 OTUs for expected_*rpoB* and expected_*16S*, respectively).

We refined the comparison between the two markers, by aggregating and aligning the nucleotide sequences obtained by Illumina-sequencing of the three replicates for each mock community and building phylogenetic trees with the maximum likelihood method. The results for the mock1 community analysis are shown (see Figure 3 and Additional file 3 for *rpoB* marker; Figure 4 and Additional file 4 for 16S marker). As a control, we built a phylogenetic tree with the sequences obtained by Sanger sequencing of both the *rpoB* and 16S markers after PCR amplification from the DNA of each separate bacterial species (Figure 3A and 4A). With the 0.1% cutoff, a comparison with the topology of the Expected_*rpoB* and Expected_16S mock communities showed aberrant clusters in Illumina-sequenced mock communities (in gray in Figures 3 and 4). We removed these clusters, which probably corresponded to chimeric sequences that had escaped the filtering process. With the *rpoB* marker, we were able to identify the 19 bacterial taxa of the mock1 community with perfect-match taxonomic identities (sensitivity = 100%), but we noted the presence of a few sequence variants (12 and 4 sequences variants in the FROGS and DADA2 analyses, respectively). For the 16S rRNA gene marker, the sensitivity for OTU detection was 100% for FROGS and 76% for DADA2, but we observed many more sequence variants than with the *rpoB* marker (24 and 25 sequence variants with FROGS and DADA2, respectively), potentially due to PCR/sequencing errors or intragenomic polymorphisms for 16S rRNA copy number. With the 1% cutoff, no chimeric sequences or sequence variants were detected with either the *rpoB* or the 16S rRNA gene marker, but the OTU detection sensitivity decreased considerably (between 58% and 64%) (Figures 3 and 4 and Additional file 3 and 4). We therefore selected the 0.1% cutoff as yielding the optimal sensitivity for OTU detection, despite the slight overestimation observed.

**Figure 3.**
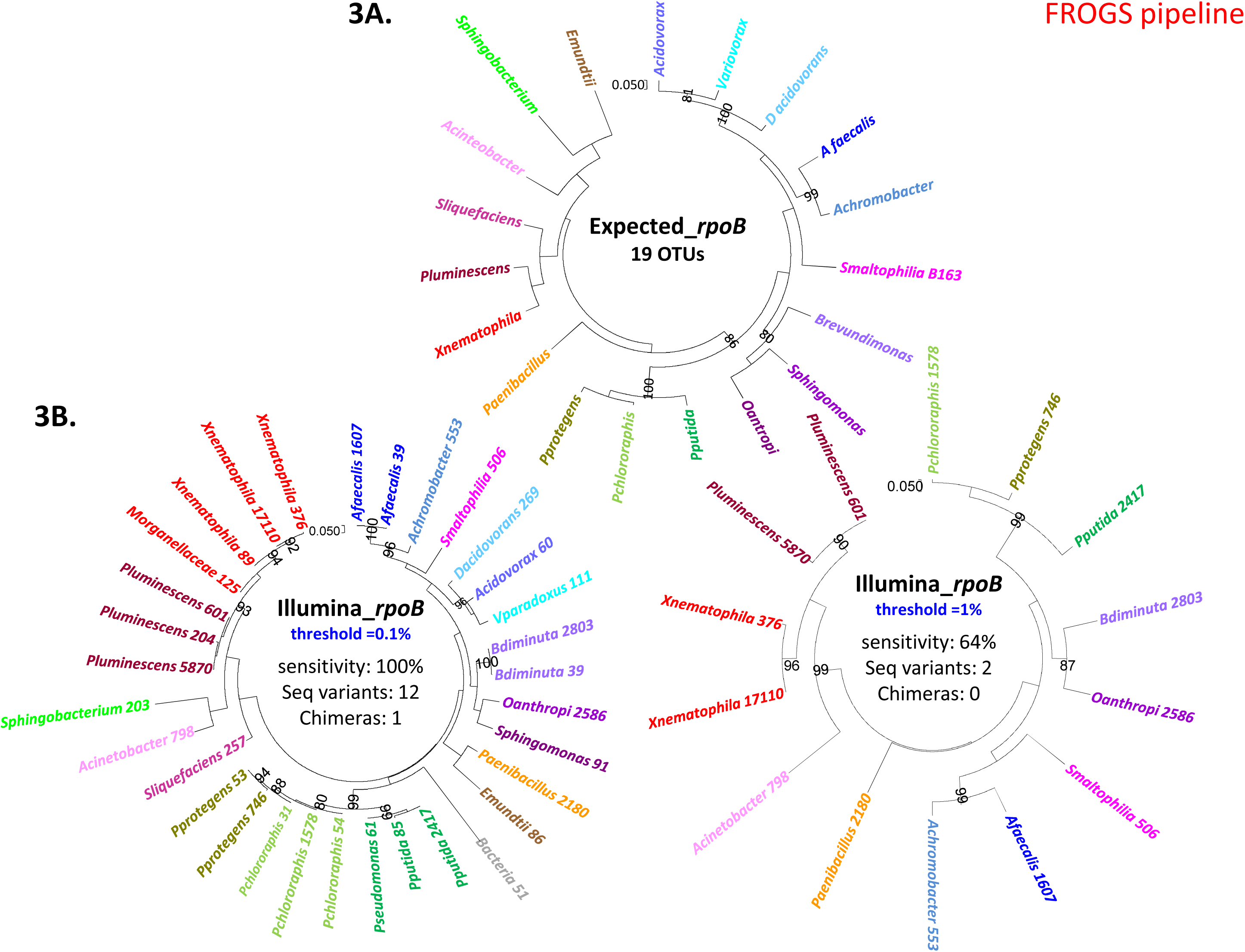
Comparison of the expected bacterial composition and the observed OTU composition obtained with Illumina-amplicon *rpoB* sequencing in the mock1 community (FROGS process). Phylogenetic trees based on the *rpoB* 430 bp-region Muscle alignment were inferred with MEGA7, with a PhyML-based maximum likelihood algorithm and the GTR model for: **(A)** individual Sanger sequencing of *rpoB* gene fragment for the 19 taxa making up the experimental mock communities (Expected_*rpoB*); **(B)** the observed OTUs obtained after Illumina-amplicon *rpoB* sequencing of the mock1 community (OTUs of the three replicates are summed), but only OTUs with an abundance >0.1% of total reads in individual replicates were included in the analysis (threshold = 0.1%); **(C)** As in B, but only OTUs with an abundance >1% of total reads in individual replicates were included in the analysis (threshold = 1%). The OTUs corresponding to the same taxa from the 19 bacterial components of the mock community are highlighted in the same color. The sum of read numbers is indicated after the OTU name. Bootstrap values (percentages of 1000 replicates) of more than 80% are shown at the nodes.

**Figure 4.**
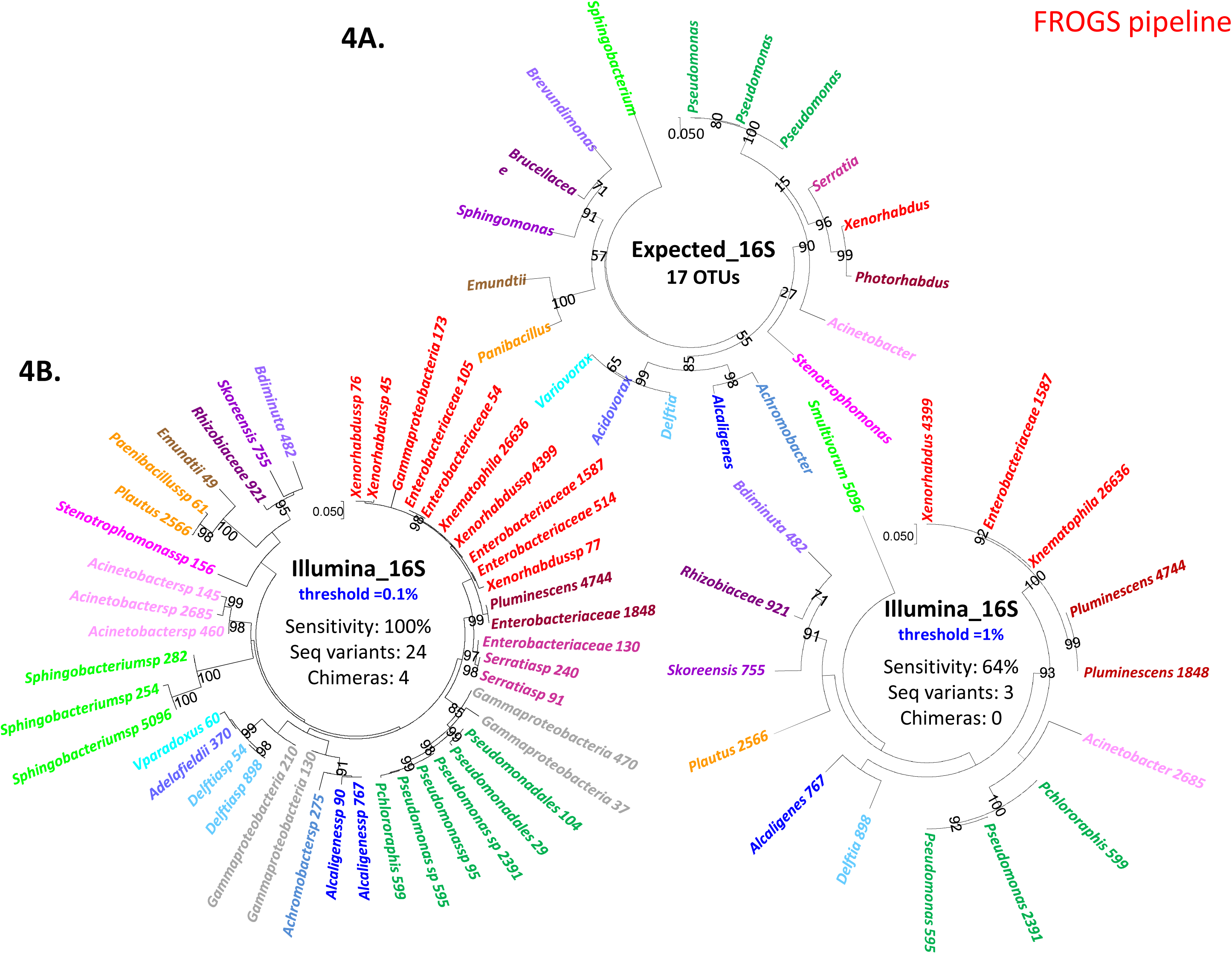
Comparison of expected bacterial composition and the observed OTU composition generated by Illumina-amplicon 16S sequencing for the mock1 community (FROGS process). Phylogenetic trees based on the 16S rRNA-V3V4 region Muscle alignment were inferred with MEGA7, a PhyML-based maximum likelihood algorithm and the GTR model for: **(A)** individual Sanger sequences of the V3V4 region of the 16S rRNA gene of the 19 taxa making up the experimental mock communities (Expected_16S); **(B)** the observed OTUs obtained after Illumina-amplicon 16S sequencing of the mock1 community (OTUs of the three replicates are summed), but only OTUs with an abundance >0.1% of total reads in individual replicates were included in the analysis (threshold = 0.1%); **(C)** As in B, but only OTUs with an abundance >1% of total reads in individual replicates were included in the analysis (threshold = 1%). The OTUs corresponding to the same taxa from the 19 bacterial components of the mock community are highlighted in the same color. The sum of read numbers is indicated after the OTU name. Bootstrap values (percentages of 1000 replicates) of more than 80% are shown at the nodes.

### Assessment of the quantitative potential of the *rpoB* and 16S markers for metabarcoding approaches

For visualization of the potential abundance biases during metabarcoding, we generated boxplots of the relative abundances of the 19 taxa making up the mock communities (as shown in Figure 5 for the Mock3 community and in Additional file 5 for the other mock communities, except for mock2, for which the highest abundance of *Xenorhabdus* and *Photorhabdus* masked the presence of other bacteria). For both *rpoB* and 16S markers, we observed differences in the relative abundances of OTUs between results from metabarcoding analysis and the expected OTU composition. For the five mock communities, we calculated the abundance bias ratios for each of the 19 taxa (Table 3). Interestingly, these ratios were not correlated with a particular phylogenetic group, and instead depended on both the bacterial strains and markers used for metabarcoding. These results confirm that, even with a single-copy gene such as *rpoB*, the measurement of OTU relative abundance is not strictly reliable, and metabarcoding should therefore be considered a semi-quantitative method.

**Table 3.** Abundance bias ratios calculated for the 19 taxa after Illumina-amplicon sequencing of rpoB and 16S rRNA for the five mock communities

|  | 16S<br>copy<br>number | Genom<br>e size<br>(Mb) | Ratio of metabarcoding relative abundance of OTUs / expected relative abundance |  |  |  |  |  |  |  |
| --- | --- | --- | --- | --- | --- | --- | --- | --- | --- | --- |
|  |  |  | Mock1 |  | Mock3 |  | Mock4 |  | Mock5 |  |
|  |  |  | *rpoB* | 16S | *rpoB* | 16S | *rpoB* | 16S | *rpoB* | 16S |
| *E. mundtii* | 6 | 3.3 | 2.21 | 0.91 | 3.44 | 1.45 | 3.61 | 1.15 | 4.21 | 1.61 |
| *D. acidovorans* | 5 | 6.8 | 0.46 | 1.42 | 0.66 | 1.86 | 0.75 | 2.03 | 0.82 | 1.93 |
| *A. xyloxisidans* | 3 | 6.8 | 0.93 | 0.68 | 1.32 | 0.97 | 1.35 | 0.94 | 1.42 | 0.85 |
| *S. liquefaciens* | 7 | 5.3 | 0.43 | 0.34 | 0.71 | 0.82 | 0.79 | 0.91 | 0.87 | 0.97 |
| *A. faecalis* | 3 | 4.2 | 2.52 | 2.37 | 3.29 | 3.15 | 3.58 | 3.34 | 3.85 | 3.31 |
| *P. protegens* | 5 | 7.0 | 0.94 | 0.60 | 1.27 | 0.85 | 1.25 | 0.87 | 1.30 | 0.88 |
| *O. anthropi* | 4 | 5.2 | 2.89 | 1.22 | 3.33 | 1.59 | 3.67 | 1.64 | 3.64 | 1.68 |
| *Acidovorax sp* | 3 | 4.9 | 0.06 | 0.58 | 0.12 | 0.86 | 0.14 | 1.00 | 0.16 | 0.98 |
| *V. paradoxus* | 2 | 7.5 | 0.12 | 0.14 | 0.11 | 0.13 | 0.14 | 0.17 | 0.07 | 0.12 |
| *P. chlororaphis* | 5 | 7.3 | 1.29 | 0.40 | 1.61 | 0.57 | 1.72 | 0.56 | 1.74 | 0.60 |
| *P. putida* | 7 | 6.2 | 1.97 | 1.42 | 1.52 | 0.92 | 1.62 | 1.14 | 1.71 | 1.07 |
| *Paenibacillus sp* | 10 | 6.0 | 1.68 | 0.84 | 1.99 | 1.25 | 2.23 | 1.17 | 2.21 | 1.28 |
| *Sphingomonas sp* | 2 | 4.1 | 0.07 | 1.26 | 0.09 | 1.75 | 0.09 | 1.90 | 0.08 | 1.78 |
| *Sphingobacterium sp* | 7 | 6.9 | 0.09 | 1.40 | 0.14 | 1.68 | 0.19 | 1.62 | 0.18 | 1.65 |
| *Brevundimonas sp.* | 2 | 3.4 | 1.13 | 0.41 | 1.21 | 0.60 | 1.30 | 0.59 | 1.25 | 0.57 |
| *Acinetobacter sp.* | 6 | 3.6 | 0.31 | 0.76 | 0.43 | 1.02 | 0.53 | 0.98 | 0.56 | 1.03 |
| *S. maltophilia* | 4 | 4.9 | 0.14 | 0.06 | 0.20 | 0.11 | 0.23 | 0.09 | 0.19 | 0.09 |
| *P. luminescens* | 7 | 5.7 | 1.56 | 0.97 | 6.66 | 4.01 | 5.60 | 2.97 | 10.76 | 3.63 |
| *X. nematophila* | 7 | 4.6 | 1.18 | 1.37 | 2.42 | 0.95 | 2.57 | 0.76 | 2.68 | 1.16 |

**Figure 5.**
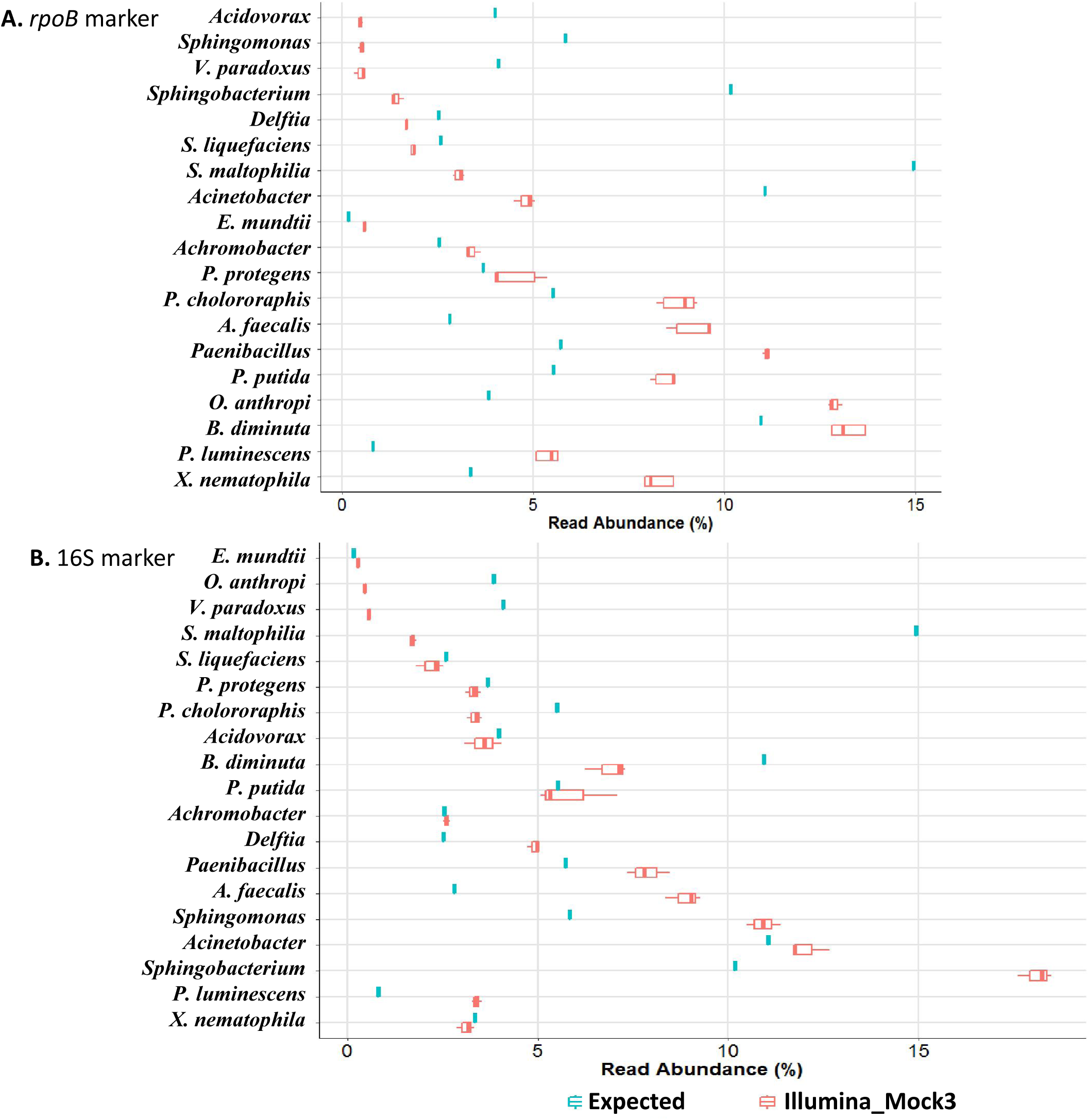
Comparison of the observed and expected relative abundances of the bacterial community of the mock3. Boxplots show the variation of OTU relative abundance in: **(A)** the mock3 community (three replicates), with use of the *rpoB* marker; (**B)** the mock3 community (three replicates) with use of the 16S marker. The blue box plots show the expected relative abundances of the species based on the theoretical composition of the mock3 community, and the red box plots show the observed relative abundance of OTUs based on Illumina-amplicon sequencing of the mock3 community (FROGS pipeline analysis). Error bars indicate the standard deviation for triplicate samples. The taxonomic identities of the 19 bacterial strains making up the mock3 community are indicated on the *y*-axis, and the relative abundance of each taxon (values = median percentage read abundance across all replicates for Illumina-sequenced mock communities) is plotted on the *x*-axis. The number of reads has been corrected with respect to the number of copies of the 16S rRNA gene in each taxon, for the calculation of observed relative abundance.

### Efficiency of the *rpoB* marker for describing the microbiota of entomopathogenic nematodes

We performed metabarcoding analyses on a nematode strain, *Steinernema glaseri* SK39 (four technical replicates per strain), with both the *rpoB* and the 16S markers as taxonomic targets. The taxonomic targets were amplified and sequenced and the sequences were processed with the FROGS and DADA2 pipelines. For the *rpoB* marker, we obtained 48,719 reads (means of 12,180 reads per sample) and 56,699 reads (mean of 14,175 reads per sample) after FROGS and DADA2 processing, respectively (Additional file 6). For 16S rRNA gene marker, we obtained 47,298 reads (mean of 11,824 reads per sample) and 41,655 reads (mean of 10,414 reads per sample) after FROGS and DADA2 processing, respectively (Additional file 6). Rarefaction curves obtained from quality-filtered reads indicated that sequencing depth was sufficient for both the 16S and *rpoB* markers (Additional file 2). We checked that the biological sample sequences were not contaminated with sequences present in the negative-control samples (Additional file 7). A comparison of the taxonomic affiliation levels of the observed OTUs again confirmed the higher taxonomic discrimination potential of the *rpoB* marker than of the 16S marker (Figure 1B). For each replicate, OTU numbers after of the application of an abundance threshold of 0.1% (for individual samples) are detailed in Additional file 8. Depending on the taxonomic marker (*rpoB* or 16S) and analysis pipeline (FROGS or DADA2) used, the number of OTUs detected ranged from 30 to 55 (Figure 6). We then analyzed the taxonomic composition of the bacterial communities present in the nematode microbiota. Similar bacterial compositions were obtained with both markers at the phylum level (Figure 7A and 7B), with the Proteobacteria the most abundant bacterial phylum. At the family level, the results differed between the two markers (Figure 7C and 7D). We refined the comparison of the markers at more discriminant taxonomic ranks, by building phylogenetic trees from OTU sequences (Figure 8 for FROGS datasets and Additional file 8 for DADA2 dataset). The *rpoB* marker accurately detected the symbiotic bacterium *X. poinarii*. By contrast, the 16S marker predicted the presence of other species from the genera *Xenorhabdus* and *Photorhabdus* (phylogenetically close species), leading to the erroneous detection of bacteria that are never associated with this nematode (Figure 8 and Additional file 8); the symbiotic bacterium *X. poinarii* was identified only by the DADA2 analysis (Additional file 8). Moreover, numerous sequence variants were detected with the 16S marker (46 and 40 sequence variants with the FROGS and DADA2 pipelines, respectively), whereas far fewer variants were detected with the *rpoB* marker (13 and 12 sequence variants with FROGS and DADA2, respectively). We also observed a few OTUs corresponding to chimeras for both markers in DADA2 analysis (Additional file 8). Once these chimeras and sequence variants were removed, the number of OTUs detected in the nematode microbiota was relatively similar between the 16S and *rpoB* markers, at about 25 OTUs (Figure 8 and Additional file 8).

**Figure 6.**
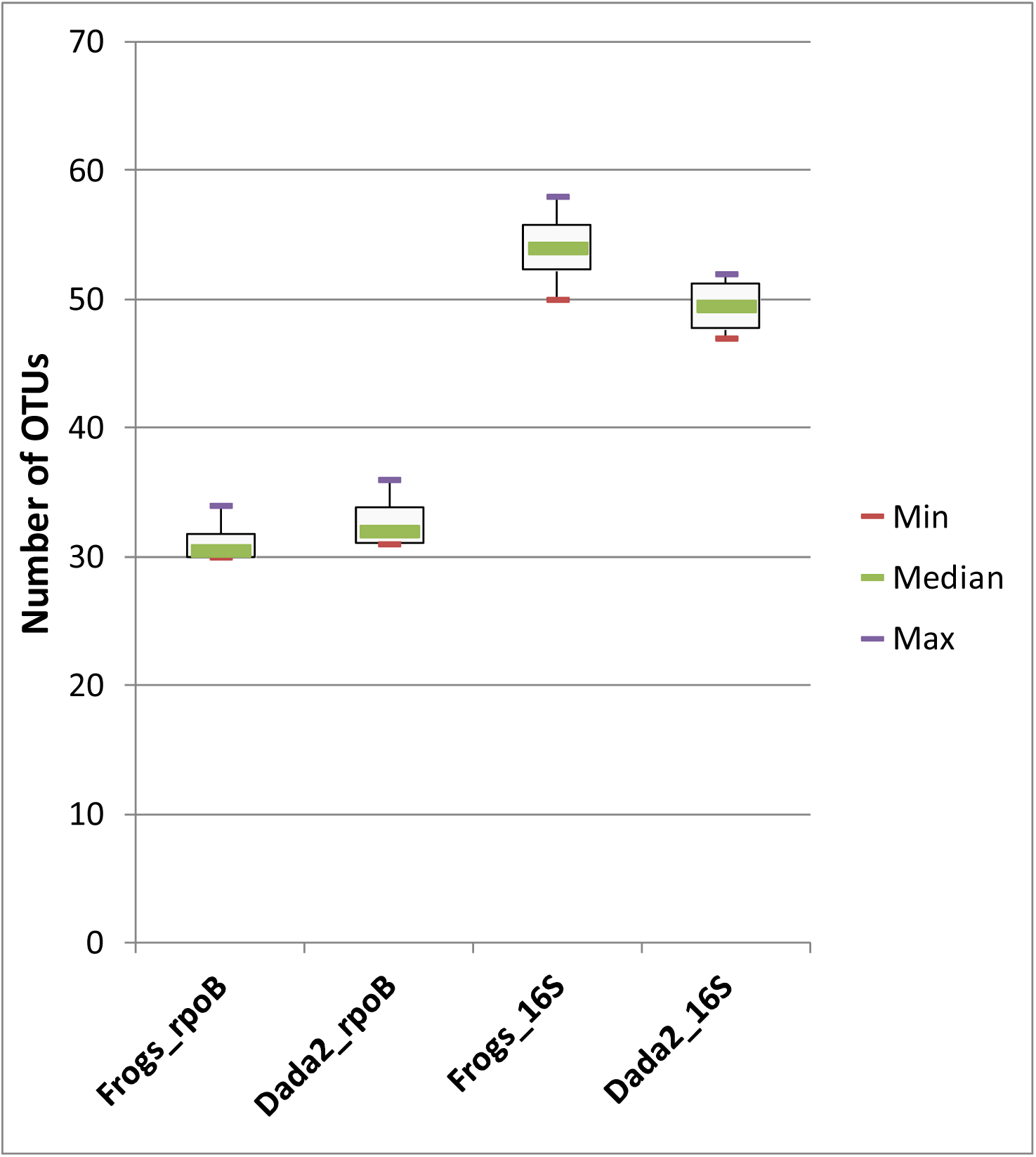
Impact of the markers and sequence analysis pipeline tool used on observed OTU richness after Illumina-amplicon sequencing of the nematode *Steinernema glaseri*. Boxplots representing the variation in OTU numbers (only OTUs with an abundance >0.1% of total reads in individual replicates) as a function of the marker (*rpoB* marker *versus* 16S marker) and sequence analyse pipeline tool (FROGS *versus* DADA2) used. Each boxplot corresponds to a statistical analysis of the four replicates; thin purple and red lines correspond to minimum and maximum values, respectively; and thicker green lines in the boxes correspond to the medians.

**Figure 7.**
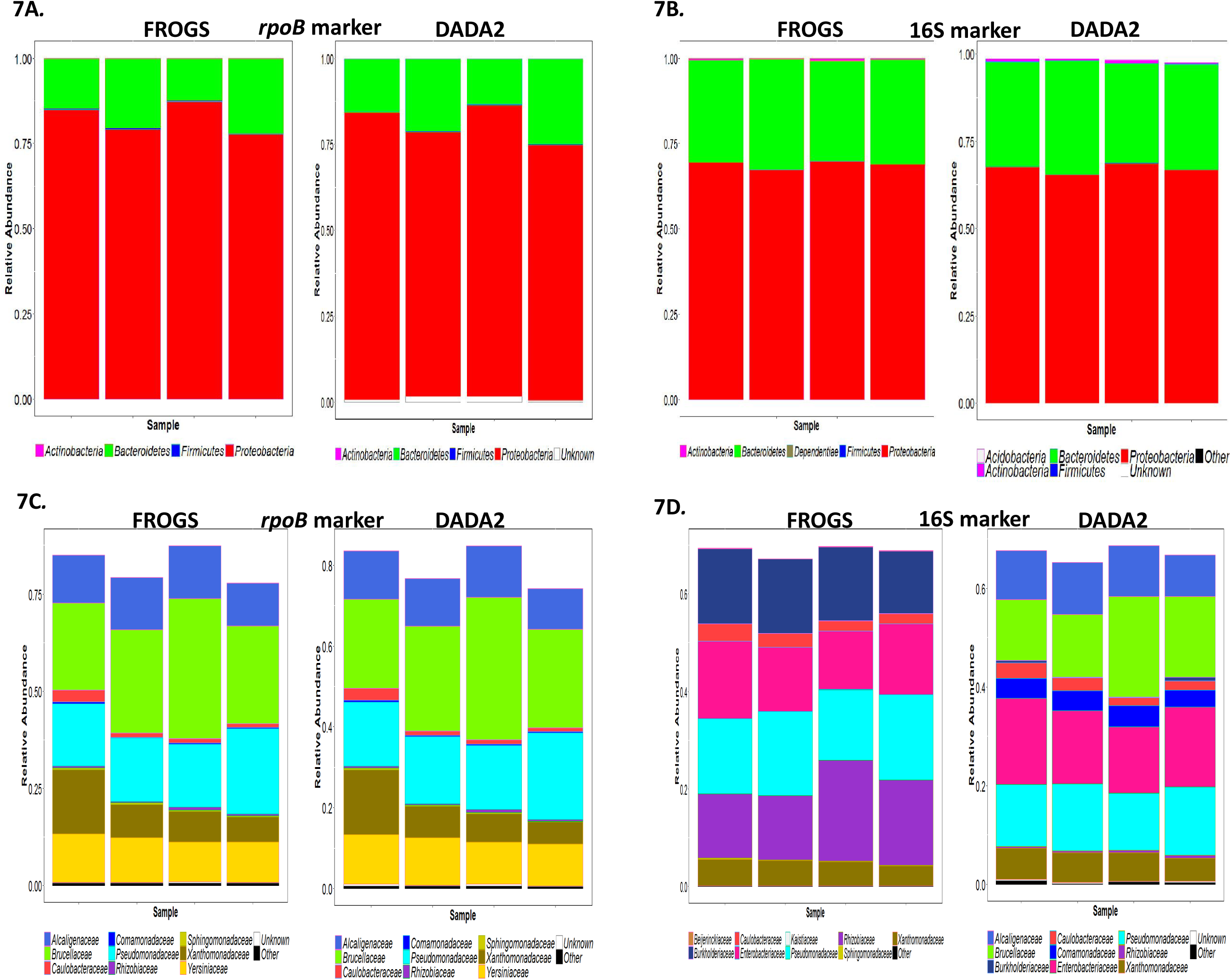
Composition plots (phylum and family levels) of the relative abundances of OTUs obtained by Illumina-amplicon sequencing of the nematode *Steinernema glaseri* SK39 with primers for *rpoB* and 16S markers. Bar plots, each representing an individual replicate, showing the relative abundance of OTUs (the data have been normalized as a % of total OTUs): (**A**) within Bacteria, at the phylum level, after amplification with *rpoB* marker; (**B**) within Bacteria, at the phylum level, after amplification with 16S marker; **(C)** within Proteobacteria, at family level, after amplification with *rpoB* marker; **(D)** within Proteobacteria, at family level, after amplification with 16S marker. Sequence datasets were processed with either the FROGS or the DADA2 pipeline, as indicated at the top of the panels (subpanels within A-D). Within each panel, the ratio of each taxon was estimated from the sum of all taxa (within all phyla or within Proteobacteria).

**Figure 8.**
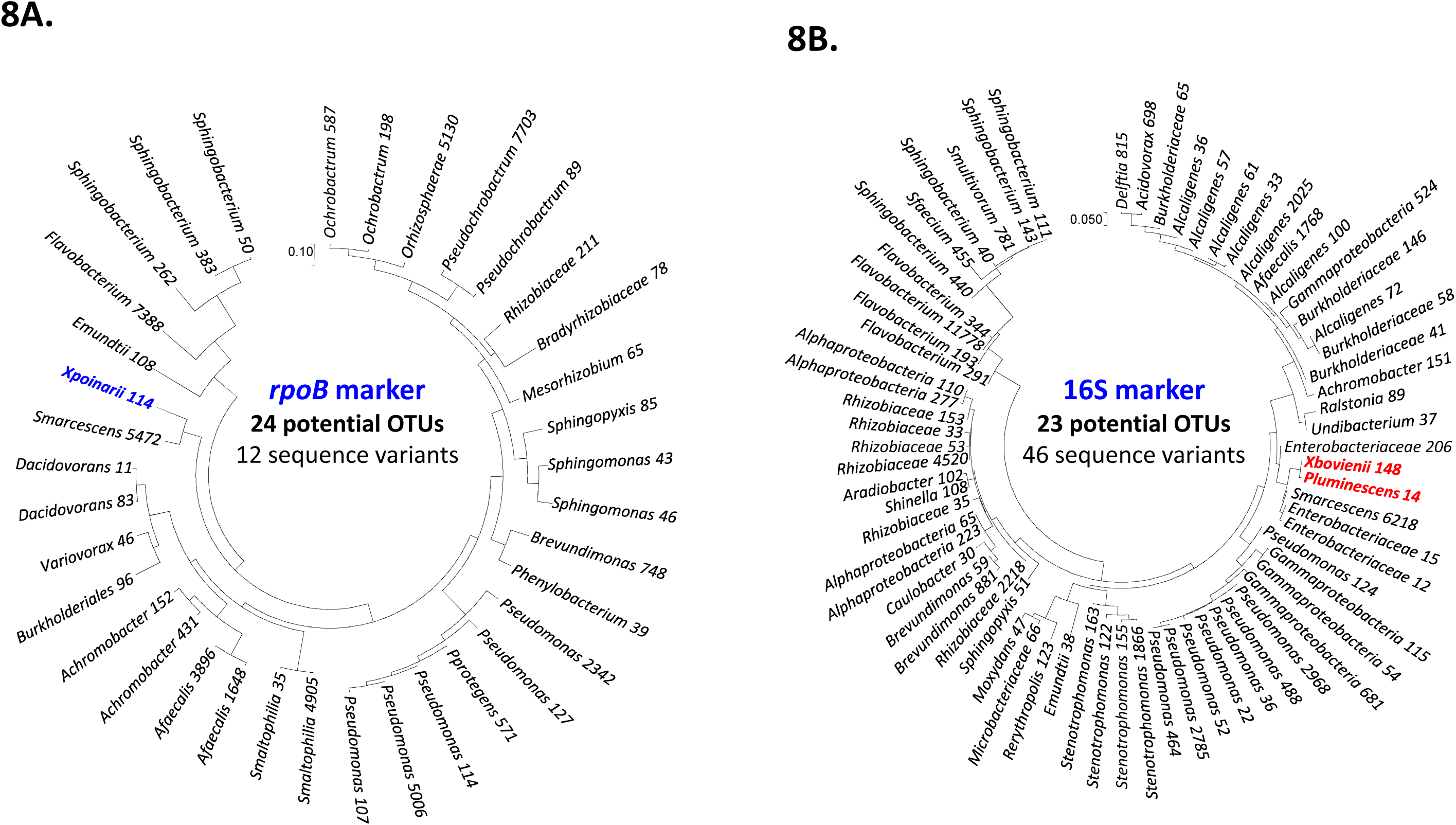
Comparison of the bacterial composition data generated by Illumina sequencing of the nematode *Steinernema glaseri* SK39 (FROGS process) with the *rpoB* and 16S markers. Phylogenetic trees were inferred with MEGA7, using a PhyML-based maximum likelihood algorithm and the GTR model: (**A)** Phylogenetic tree of the observed OTUs obtained after Illumina-amplicon *rpoB* sequencing; (**B)** Phylogenetic tree of the observed OTUs obtained after Illumina-amplicon 16S rRNA sequencing. The OTUs of the three replicates are summed, but only OTUs with an abundance >0.1% of total reads in individual replicates were included in the analysis. The OTUs corresponding to the true symbiotic bacterium (*X. poinarii*) are highlighted in blue. The OTUs corresponding to false-positive symbiotic bacteria from the genera *Xenorhabdus* and *Photorhabdus* are highlighted in red. The sum of the read numbers is indicated after the OTU name. Bootstrap values (percentages of 1000 replicates) of more than 80% are shown at the nodes.

## Discussion

Metabarcoding methods, generally based on the high-throughput sequencing of 16S amplicons, are widely used. Despite the known bias of “universal” 16S markers, only a few studies have reported the use of alternative markers. We assessed the potential benefit of a portion of the *rpoB* gene as an alternative genetic marker.

We analyzed the sequence data generated by metabarcoding with *rpoB* and 16S markers on an artificial bacterial DNA complex corresponding to 19 different phylogenetic taxa. One key factor determining the choice of taxonomic markers for metabarcoding studies is the ability of the marker to distinguish between OTUs at the lowest possible taxonomic rank. We found that taxonomic assignation was more accurate with the housekeeping gene *rpoB* than with the 16S marker. The *rpoB* gene classified OTUs more effectively to species level, with the resolution of the 16S marker frequently limited to the genus or a higher taxonomic level. It can be difficult to resolve the taxonomy of 16S rRNA gene sequences based on a limited segment, such as the V3V4 region. Closely related bacteria, such as those of the Enterobacteriaceae family, cannot be differentiated solely on the basis of differences in the V3V4 region [23]. The composition and abundance of OTUs in mock communities are known *a priori*. Such mock communities are therefore useful tools for detecting potential biases during method development and for optimizing data analysis pipelines [24].

We show here that the *rpoB* marker gave results that more closely matched the expected composition of the mock community. When the 16S marker was used, we observed a strong distortion of the composition of the bacterial community obtained by the metabarcoding of the mock communities away from the expected bacterial composition, with the values obtained for OTU richness composition higher than the actual OTU richness. This OTU overestimation bias was weaker for the *rpoB* marker. By contrast to the single-copy housekeeping gene targeted by the *rpoB* marker, sequence heterogeneity between the different copies of the16S marker may lead to the amplification of numerous sequence variants during metabarcoding, resulting in the identification of excessive numbers of OTUs in 16S datasets. Intragenomic ribosomal diversity [25] or ribosomal paralogs [26] have frequently been implicated as the major source of sequence variants in 16S Illumina amplicon-sequencing analyses. The observed OTU inflation may also be explained by cumulative errors occurring during the amplification and sequencing steps of the metabarcoding procedure, resulting in the detection of sequence variants. Interestingly, given that *rpoB* is a protein-coding gene, sequence errors can be readily identified and removed if they disrupt the reading frame [12], providing an added benefit of housekeeping genes as targets in metabarcoding studies. Some authors have also suggested that excessive OTU diversity may be at least partially explained by the presence of unfiltered chimeric reads [27], and laboratory contaminants [28]. Nelson and collaborators reported a strong overestimation of mock community diversity (25-125 times higher than expected) in the absence of careful checking of the data. Similarly, Kunin and collaborators found that diversity was grossly overestimated for their mock community data unless a quality threshold was implemented. We show here that the removal of OTUs with read abundances below 1% (for individual samples) decreases the number of sequence variants. However, this threshold cutoff of 1% is much less sensitive than the 0.1% threshold cutoff for describing the bacterial communities making up the mock communities.

When using metabarcoding for quantitative analyses, caution is required concerning the conclusions drawn about the relative abundances of bacterial taxa, even with a single-copy gene, such as *rpoB*. Our results highlight the existence of a bias in abundance taxa, this bias being strain-dependent and varying with the marker used. This bias probably reflects the amplification bias occurring during PCR cycles [6].

We finally used the *rpoB* marker to analyze microbial communities in an entomopathogenic nematode, *Steinernema glaseri*, known to carry an intestinal symbiotic bacterium, *Xenorhabdus poinarii,* but for which microbiota composition remains otherwise unknown. We found that the findings concerning the bacterial community associated with *S. glaseri* depended on the marker used. The *rpoB* marker gave better taxonomic discrimination and was capable of reliably identifying the symbiotic bacterium. The 16S rRNA marker is able to detect the symbiont only with one of the pipelines, as well as false positive OTUs phylogenetically related to the symbiont.

The *rpoB* marker was successfully used here to describe the microbiota of an entomopathogenic nematode, with a microbiota of low diversity including bacterial taxa for which *rpoB* sequences are routinely available from databases. Further testing of the *rpoB* marker is required in other types of complex microbiota, such as those containing members of phyla other than Firmicutes and Proteobacteria. Despite its disadvantages, the 16S marker should not be entirely abandoned, particularly for the exploration of complex and unknown ecological niches, as it is a reference marker for which a very rich and complete database is available for taxonomic assignment. For high-throughput amplicon sequencing studies, we therefore recommend the use of multigenic approaches based on different taxonomic markers, targeting at least one housekeeping gene (*rpoB* is a good candidate) and a variable16S rRNA gene region.

## Conclusions

The use of 16S sequencing raises a number of challenges, including primer bias, gene copy number, PCR or sequencing artifacts and contamination. Metabarcoding approaches, which target a taxonomically relevant marker, such as the *rpoB* gene, are a potential alternative, making it possible to overcome at least some of these challenges. The major benefit of *rpoB* sequencing is its potential for improving taxonomic assignment and for more detailed investigations of OTU richness at species level, providing a more accurate description of the composition of microbiota communities. The greater ability of *rpoB*-based analyses to discriminate between phylogenetically different groups of species should increase resolution and provide more reliable results for metagenomic studies. Our results also highlight the need to develop and use mock community as controls for all microbial studies, to pick up potential sequence errors, which may arise at any step in next-generation sequencing protocols. For Illumina amplicon-sequencing strategies, we also strongly recommended the use of a multigenic approach based on at least two taxonomic markers, including a protein-coding gene, such as *rpoB,* and the 16S rRNA marker.

## Methods

### Biological material

The nematode/bacterial strains and primers used in this study are listed in Additional file 9.

### Isolation, multiplication and storage of nematode strains

*Steinernema* nematodes were originally isolated with an *ex situ Galleria* trap, as previously described [30]. The nematodes used here had been stored for decades in the DGIMI collection. Infective juveniles (IJs) were stored in Ringer’s solution (Merck) and were multiplied every six months by infestation of the last instar of *Galleria*. Briefly, IJs were added to *Galleria* larvae in Petri dishes (laboratory *Galleria* trap). When the *Galleria* larvae died, their cadavers were placed on a white trap and the IJs that emerged from them were stored in Ringer’s solution at 9°C.

### Isolation, multiplication and storage of the bacteria used in the mock communities

The symbiotic bacteria, *Xenorhabdus* and *Photorhabdus,* were isolated by the hanging drop technique [31]. Bacteria from other genera were isolated from crushed IJs nematodes or from the contents of *G. mellonella* cadavers after nematode infestation. For the isolation of bacteria from entomopathogenic nematodes, 20 IJs were placed in a 1.5 mL Eppendorf tube containing 200 µL of Lysogeny Broth medium and three 3-mm glass beads, and were subjected to three cycles of grinding (1 minute, at 30 Hz followed by 1 minute without agitation) in a TissueLyser II apparatus (Qiagen, France). The solutions obtained from crushed nematodes or the contents of *G. melonella* were plated on both nutrient agar (Difco) plates and nutrient bromothymol blue agar (NBTA) plates [32], and incubated at 28°C for 48 hours. Bacterial colonies with different morphotypes were stored at -80°C in 16% glycerol (v/v).

### Molecular identification of the bacteria isolated and mock community design

Bacterial isolates were identified as previously described [33], by amplifying and sequencing a near full-length 16S rRNA gene (1372 bases). Briefly, bacterial genomic DNA was extracted as previously described [34] and stored at 4°C. The 16S rRNA gene was amplified (the primer sequences are indicated in Additional file 9) in a Bio-Rad thermocycler (Bio-Rad, USA). PCR products were analyzed by agarose gel electrophoresis. Sanger sequencing of the 16S rRNA amplicons was performed by MWG-Eurofins (Deutschland), and sequences were blasted against the NCBI database for taxonomic identification of the bacterial isolates. Nineteen bacterial isolates encompassing a broad taxonomic diversity among eubacteria were selected to compose the reference mock communities (Table 1). The DNA concentration of each selected bacterial strain was quantified on a Qubit Fluorometer (Thermo Fisher Scientific, USA) and five mock communities were generated by mixing the DNA of the 19 strains (the DNA concentrations of symbiotic taxa in the mock communities varied over four orders of magnitude). The proportions of the 19 bacterial taxa composing the five mock communities, in genome number equivalents, are shown in Table 2. The number of genome equivalents was calculated as follows: number of genome copies = [DNA amount (ng) * 6.022×10^23^] / [Genome size (bp) * 1×10^9^ * 650 g/mole of bp].

### Sanger sequencing of the *rpoB* and V3V4 regions of the 19 bacterial isolates present in the mock communities

The *rpoB* and V3V4 regions were amplified as described above (the primer sequences are indicated in Additional file 9). Sanger sequencing of the amplicons was performed by MWG-Eurofins (Deutschland).

### DNA extraction from IJs

We performed four DNA extraction replicates per nematode sample, each replicate consisting of about five thousand IJs sampled from a storage batch. We minimized the risk of contamination with microorganisms from the body surface, by washing the IJs with copious amounts of tap water on a filter. The washed IJs were recovered from the filter with a sterile pipette, transferred to 10 mL of sterile ultrapure water and immediately frozen at - 80°C for future use. DNA was extracted from the IJs with the Quick Extract kit (Epi-centre, USA). Briefly, frozen samples were rapidly thawed, heated at 80°C for 20 minutes and centrifuged (2,500 x *g*, 10 minutes) to pellet the IJs. Once the supernatant had been removed, we added 200 µL of Quick Extract lysis solution and the mixture was transferred to 2 mL Eppendorf tubes containing three sterile 3-mm glass beads. IJs were crushed by three cycles of mechanical grinding (2 minutes at 30 Hz followed by 1 minute without agitation) in a TissueLyser II apparatus (Qiagen, France). We added 2 µL of Ready-Lyse Lysozyme Solution (Epi-centre, USA) to the ground samples, which were then incubated at room temperature until the solution cleared (24 to 48 hours). Samples were again heated at 80°C for 20 minutes, and the lysis of IJs was checked under a light microscope. After complete lysis, the samples were subjected to an additional treatment with 20 µL of 20 mg/mL RNaseA (Invitrogen PureLinkTM RNaseA, France). Finally, a phenol-chloroform purification step was performed, followed by chloroform purification. The DNA was precipitated in absolute ethanol, washed twice in 70% ethanol, resuspended in 50 μL ultrapure water and stored at - 20°C. The DNA products were analyzed by agarose gel electrophoresis and quantified with a Nanodrop spectrophotometer (Thermo Fisher Scientific). We assessed the levels of contaminating bacterial DNA at each of the various steps in DNA sample preparation, by including several negative control extractions with sterile ultra-pure water.

### Design of *rpoB* primers suitable for use in MiSeq Illumina sequencing

Degenerate consensual pairs of *rpoB* primers called Univ_rpoB_F_deg (forward primer) and Univ_rpoB_R_deg (reverse primer) were manually designed from clustalW alignments (http://multalin.toulouse.inra.fr/multalin/) of *rpoB* gene sequences from bacterial reference genomes covering a broad taxonomic diversity among eubacteria. The binding sites of the selected primers correspond to *Escherichia coli* K12 nucleotide positions 1630 to 2063, making it possible to amplify a 434-nucleotide portion of the *rpoB* gene. The specificity of the *rpoB* primers was checked *in silico* on a large panel of publicly available complete sequenced genomes with the “Blast and Pattern Search” web tool implemented on the *MaGe* annotation platform (http://www.genoscope.cns.fr/agc/mage). The universality of the *rpoB* primers was checked *in silico* on a large panel of *rpoB* genomic sequences publicly available from the NCBI database, with the Primers toolbox implemented in CLC Genomics Workbench 3.6.1.

### Library preparation and Illumina MiSeq sequencing

Amplicon libraries were constructed following two rounds of PCR amplification. The first amplification step was performed with the high-fidelity iProof™ DNA Polymerase (BioRad), in a Bio-Rad thermocycler, on 10 to 100 ng of DNA. The hypervariable V3V4 region of the 16S rRNA gene was targeted with the universal primers F343 and R784, and the *rpoB* fragment was targeted with the previously designed primers Univ_rpoB_F_deg and Univ_rpoB_R_deg (the primer sequences are shown in Additional file 9). Thirty amplification cycles were performed with annealing temperatures of 65°C and 60°C for the V3V4 region (∼ 460 base pairs), and the *rpoB* region (∼ 435 base pairs), respectively. We assessed the amount of contaminating DNA in these PCRs, by including negative PCR controls with sterile ultra-pure water as the template. The amounts of amplicon DNA and amplicon sizes were analyzed by agarose gel electrophoresis. Single multiplexing was performed at the Genomics and Transcriptomics Platform (INRA, Toulouse, France), with 6 base pairs index sequences, which were added to reverse primers during a second PCR with 12 cycles. Amplicon libraries were sequenced with Illumina MiSeq technology (MiSeq Reagent Kit v2) according to the manufacturer’s instructions. FastQ files were generated at the end of the run. The FastQC program [35] was used for quality control on raw sequence data and the quality of the run was checked internally, by adding 30% PhiX sequences at a concentration of 12.5 pM. Each paired-end sequence was assigned to its sample with the previously integrated index, and paired-end reads were assembled with FLASH [36]. Unassembled reads were discarded. The raw sequence data can be downloaded from http://www.ebi.ac.uk/ena/PRJEB24936 (accession numbers: ERS2212114-ERS2212143 for the mock communities; ERS2715153-ERS2715156 and ERS2715252-ERS2715255 for the nematode samples; ERS2715166-ERS2715174 and ERS2715260-ERS2715268 for the extraction-control samples)

### rpoB sequence database design

For taxonomic assignment with the *rpoB* marker, we constructed a reference database including ∼ 45000 sequences; this database is available from the FROGS website (http://frogs.toulouse.inra.fr/).

The *rpoB* sequences were collected and the quality of the resulting database was checked as previously described [9] for the *gyrB* database. Briefly, *rpoB* sequences were retrieved from 47,175 genomes sequences publicly available from the IMG database [37] at the time of analysis. CDSs belonging exclusively to the TIGR02013 protein family were defined as RpoB homologs and were retrieved for further analysis (approximately 46,300 hits found in 45,500 genomic sequences). The corresponding nucleotide sequences of the selected region used for Illumina sequencing (434 nucleotides - see above) were aligned as previously described [9]; redundant and aberrant sequences (sequences containing ambiguous nucleotides or with sequences of less than 430 nucleotides) were removed from the database.

### Bioinformatic processing of sequence data

The sequence reads obtained were processed according to the FROGS (Find Rapidly OTUs with Galaxy Solution) pipeline [38] and the DADA2 pipeline [39].

### The FROGS process

A preprocessing tool was used to remove sequences not bound to both primers, to trim the primers, and to remove all sequences containing an ambiguous base. Sequence clustering was performed with the Swarm algorithm [40]. Chimera sequences were detected with the VSEARCH algorithm, by the *de novo* UCHIME method [41, 42], and were removed. A filtering tool was used to remove spurious clusters, with read abundances of less than 0.005% of the total number of reads. The filtered sequences were assigned to taxa with RDP Classifier [43] and the 16S rRNA database SILVA pintail quality 100% [3] for V3V4 reads and the *rpoB* database for *rpoB* reads. The sequences were clustered into OTUs with a 97% similarity cutoff (with a bootstrap confidence of 80%). The OTU abundance tables are available in Additional file 10.

### The DADA2 process

The DADA2 method was developed for the analysis of short-read amplicon sequences [39]. The pipeline is based on a complete bioinformatic workflow including quality filtering, dereplication, sample inference, chimera removal, and optionally, a taxonomic assignment step. The DADA2 software takes raw amplicon sequencing data in fastq files as input, and produces an error-corrected table of the abundances of amplicon sequence variants in each sample (an ASV table) as output. As for the FROGS process, the sequence variants were assigned to taxa with RDP Classifier (sequence similarity threshold = 97%, bootstrap confidence cutoff = 80%) and the 16S rDNA database SILVA [3] for V3V4 reads and the *rpoB* database for *rpoB* reads. The OTU abundance tables are available in Additional file 10.

### Bacterial community and statistical analyses

OTU diversity and statistical analyses were performed with the R packages Phyloseq [44], Vegan [45], and Ampvis 2 (https://github.com/MadsAlbertsen/ampvis2). Briefly, rarefaction curves were calculated with Phyloseq R packages. Beta diversity was analysed with custom-developed Phyloseq command lines. A PCoA analysis (Ampvis 2 R package) based on the Bray-Curtis dissimilarity matrix was used to visualize the differences between the microbial communities of the nematodes and the microbial contaminants associated with the extraction of control samples. The significance of the clustering on PCoA plots was assessed by multivariate PERMANOVA in the Phyloseq R package on a Bray-Curtis similarity matrix with a type III of sum of squares, 9999 permutations and unrestricted permutations of raw data. The amp_boxplot function (Ampvis2 R package) was used to generate boxplots of the OTU relative abundances. All the R plots in the study were generated with the ggplot2 R package [46].

### Phylogenetic analysis

Phylogenetic analysis of the V3V4 and *rpoB* amplicons, sequence alignment and maximum likelihood analysis with the Generalised time-reversible (GTR) model were performed as described elsewhere [22]. The Mega7 tool [47] was used to generate circular phylogenetic trees.

## Supporting information

Sequence reads and OTU numbers obtained by Illumina-amplicon sequencing for rpoB and 16S markers in 15 mock community samples

Rarefaction curves obtained by Illumina-amplicon sequencing of rpoB (A) and 16S (B) markers in 15 mock community samples

Comparison of the expected bacterial composition and the observed OTU composition generated by Illumina-amplicon rpoB sequencing for the mock1

Comparison of expected bacterial composition and the observed OTU composition obtained by Illumina-amplicon 16S rRNA gene sequencing for the mock1

Comparison of the observed and expected relative abundances of the bacterial communities obtained by Illumina-amplicon rpoB(A) and 16S(B) sequencing

Sequence reads and OTU numbers obtained by Illumina-amplicon sequencing of rpoB and 16S markers for the nematode samples (four replicates).

Comparison of the bacterial communities associated with nematode samples (Steinernema glaseri SK39) and extraction control samples.

Comparison of the bacterial compositions obtained by Illumina sequencing of rpoB and 16S rRNA for the nematode S. glaseri SK39 (DADA2 process).

List of biological materials used in the experimental study

OTU abundance tables after Illumina-amplicon sequencing of rpoB or 16S markers and use of the FROGS or DADA2 sequence analysis pipeline.

## List of abbreviations

BLAST: Basic Local Alignment Search Tool
NCBI: National Center for Biotechnology Information
EMBL: European Molecular Biology Laboratory
FROGS: Find Rapidly OTUs with Galaxy Solution
OTU: Operational taxonomic unit
PCR: Polymerase chain reaction
RDP Classifier: Ribosomal Database Project (RDP) Classifier
IJs: Infective Juveniles
GTR model: Generalised time-reversible model
NBTA: nutrient bromothymol blue agar
DNA: Deoxyribonucleic acid

## Declarations

### Ethics approval and consent to participate

Not applicable

### Consent for publication

Not applicable

### Availability of data and material

The datasets generated and analyzed in this study are available from the ENA (European Nucleotide Archive) repository (http://www.ebi.ac.uk/ena/data/view/PRJEB24936)

### Authors’ contributions

JCO and SG conceived the study. JCO and SP designed and performed the experiments. JCO, SP, MG, MB and SG analyzed the data. JCO and SG supervised the project. All authors wrote, read and approved the manuscript.

## Acknowledgments

The authors thank Alain Givaudan and Julien Brillard for the proofreading of the manuscript.

## Additional files

### Additional file 1

Sequence reads and OTU numbers obtained by Illumina-amplicon sequencing for *rpoB* and 16S markers in 15 mock community samples (five mock communities, three replicates per mock community)

### Additional file 2

Rarefaction curves obtained by Illumina-amplicon sequencing of *rpoB* (A) and 16S (B) markers in 15 mock community samples (five mock communities, three replicates per mock) and the four replicates of the *Steinernema glaseri* nematode sample. Rarefaction curves were assembled, with an estimation of species richness (*x*-axis), defined with a sequence identity cutoff of 97%, relative to the total number of bacterial sequences identified (*y*-axis). Samples are presented separately, with blue lines corresponding to the mock community samples and red lines corresponding to the nematode samples.

### Additional file 3

Comparison of the expected bacterial composition and the observed OTU composition generated by Illumina-amplicon *rpoB* sequencing for the mock1 community (DADA2 process). *See Figure 3 for Figure legend details.*

### Additional file 4

Comparison of expected bacterial composition and the observed OTU composition obtained by Illumina-amplicon *16S rRNA gene* sequencing for the mock1 community (DADA2 process). *See Figure 4 for Figure legend details.*

### Additional file 5

Comparison of the observed and expected relative abundances of the bacterial communities obtained by Illumina-amplicon *rpoB* (A) and 16S (B) sequencing for the mock1, mock2, and mock3 communities. *See Figure 5 for Figure legend details.*

### Additional file 6

Sequence reads and OTU numbers obtained by Illumina-amplicon sequencing of *rpoB* and 16S markers for the nematode samples (four replicates).

### Additional file 7

#### Comparison of the bacterial communities associated with nematode samples (*Steinernema glaseri* SK39) and extraction control samples

Principal coordinate analysis (PCoA) was performed based on Bray-Curtis distances between nematode and control samples after Illumina-amplicon sequencing of the **(A)** *rpoB* marker; **(B)** the 16S marker. Nematode samples are shown as blue triangles, extraction control samples are shown as pink circles. Each point represents an individual replicate. The proportion of the variance explained by each axis is indicated as a percentage. The difference between the nematode and extraction control samples sets is statistically significant for both the *rpoB* marker (Permanova, df=1, R^2^=0.81, *p*-value=0.0015) and the 16S marker (Permanova, df=1, R^2^=0.93, *p*-value=0.0015).

### Additional file 8

Comparison of the bacterial compositions obtained by Illumina sequencing of *rpoB* and 16S rRNA for the nematode *S. glaseri* SK39 (DADA2 process). *See Figure 8 for Figure legend details.*

### Additional file 9

List of biological materials used in the experimental study

### Additional file 10

OTU abundance tables after Illumina-amplicon sequencing of *rpoB* or 16S markers and use of the FROGS or DADA2 sequence analysis pipeline.

