## Supplementary material for "*rpoB*, a promising marker for analyzing the diversity of bacterial communities by amplicon sequencing": Sequence reads and OTU numbers obtained by Illumina-amplicon sequencing for rpoB and 16S markers in 15 mock community samples

**Additional file 1**

|  |  | | | | Number of OTUs | | | | |
| --- | --- | --- | --- | --- | --- | --- | --- | --- | --- |
| Sample ID | Read number after filtering | | | | Reads ≥ 0.1% | | | Reads ≥ 1% | |
|  | FROGS | | DADA2 | | FROGS | | DADA2 | FROGS | DADA2 |
| rpoB-Mock1-rep1 | 13618 | | 15290 | | 33 | | 23 | 14 | 12 |
| rpoB-Mock1-rep2 | 14068 | | 15947 | | 28 | | 23 | 13 | 12 |
| rpoB-Mock1-rep3 | 14096 | | 16034 | | 29 | | 25 | 13 | 12 |
| rpoB-Mock2-rep1 | 14924 | | 17205 | | 22 | | 19 | 8 | 8 |
| rpoB-Mock2-rep2 | 15252 | | 17720 | | 17 | | 17 | 7 | 7 |
| rpoB-Mock2-rep3 | 17694 | | 20559 | | 19 | | 16 | 8 | 8 |
| rpoB-Mock3-rep1 | 15830 | | 17829 | | 25 | | 23 | 15 | 15 |
| rpoB-Mock3-rep2 | 14006 | | 15533 | | 26 | | 22 | 15 | 15 |
| rpoB-Mock3-rep3 | 14782 | | 16648 | | 28 | | 22 | 15 | 15 |
| rpoB-Mock4-rep1 | 16320 | | 18063 | | 30 | | 23 | 13 | 14 |
| rpoB-Mock4-rep2 | 14076 | | 15504 | | 26 | | 23 | 14 | 14 |
| rpoB-Mock4-rep3 | 12504 | | 13542 | | 28 | | 23 | 13 | 14 |
| rpoB-Mock5-rep1 | 15569 | | 17229 | | 25 | | 21 | 13 | 13 |
| rpoB-Mock5-rep2 | 16380 | | 18344 | | 27 | | 22 | 13 | 13 |
| rpoB-Mock5-rep3 | 15010 | | 16516 | | 26 | | 24 | 13 | 13 |
| 16S-Mock1-rep1 | 24716 | | 17521 | | 31 | | 32 | 9 | 14 |
| 16S-Mock1-rep2 | 18620 | | 13528 | | 43 | | 36 | 15 | 20 |
| 16S-Mock1-rep3 | 18551 | | 13669 | | 39 | | 38 | 16 | 21 |
| 16S-Mock2-rep1 | 23749 | | 16390 | | 32 | | 31 | 8 | 12 |
| 16S-Mock2-rep2 | 16590 | | 11603 | | 31 | | 27 | 8 | 12 |
| 16S-Mock2-rep3 | 23510 | | 16376 | | 33 | | 32 | 8 | 13 |
| 16S-Mock3-rep1 | 12275 | | 10567 | | 49 | | 39 | 18 | 23 |
| 16S-Mock3-rep2 | 13535 | | 11431 | | 48 | | 40 | 18 | 24 |
| 16S-Mock3-rep3 | 19005 | | 15662 | | 44 | | 46 | 18 | 23 |
| 16S-Mock4-rep1 | 14794 | | 12624 | | 37 | | 37 | 15 | 20 |
| 16S-Mock4-rep2 | 15986 | | 13391 | | 39 | | 43 | 15 | 19 |
| 16S-Mock4-rep3 | 17255 | | 14214 | | 40 | | 41 | 16 | 19 |
| 16S-Mock5-rep1 | 16477 | | 13729 | | 38 | | 34 | 15 | 19 |
| 16S-Mock5-rep2 | 17523 | | 14371 | | 34 | | 36 | 15 | 19 |
| 16S-Mock5-rep3 | 13908 | | 11565 | | 36 | | 32 | 16 | 19 |
