## Supplementary figures and images for "*rpoB*, a promising marker for analyzing the diversity of bacterial communities by amplicon sequencing"

### Comparison of the bacterial communities associated with nematode samples (Steinernema glaseri SK39) and extraction control samples.

#### Slide 1
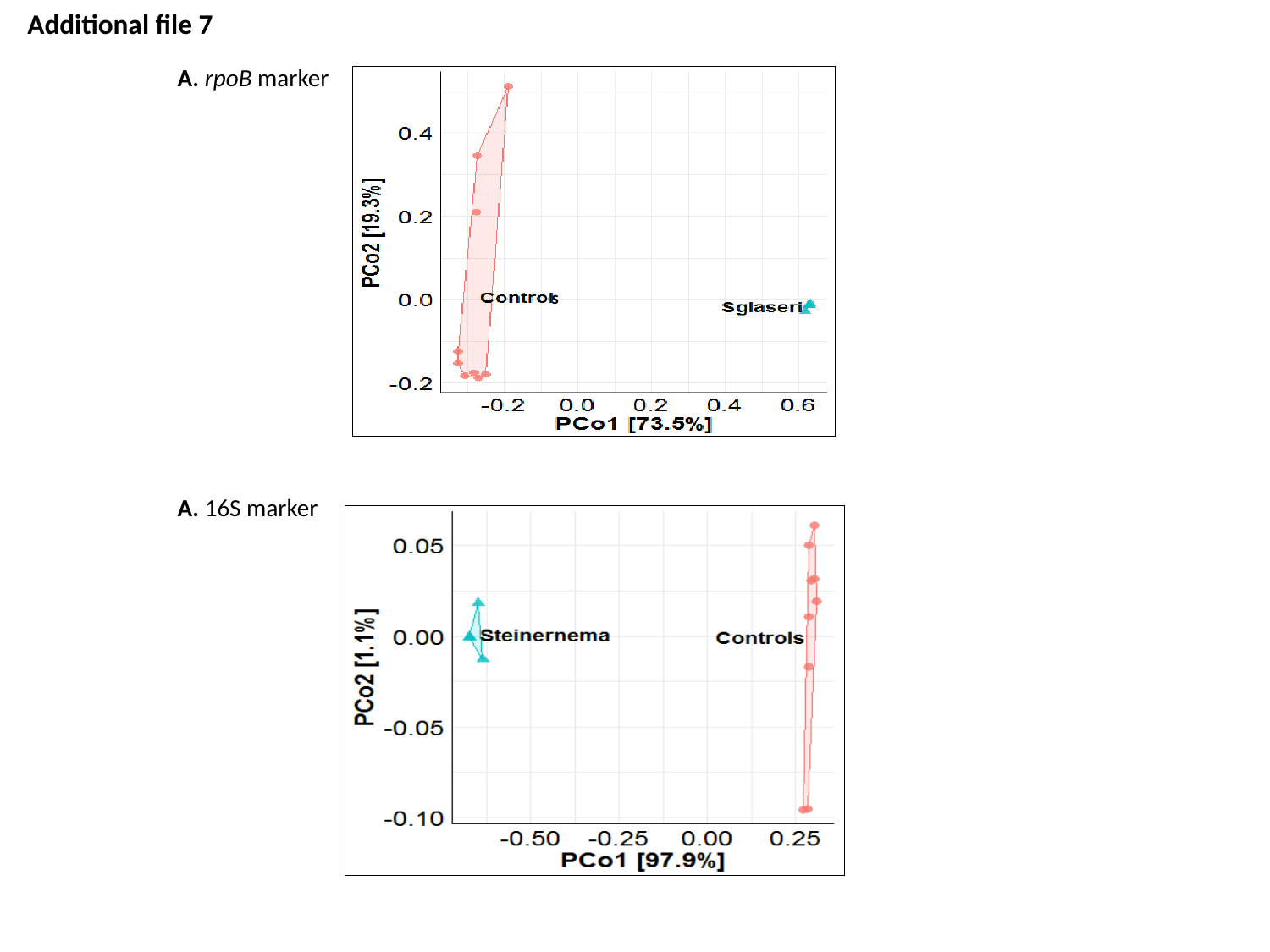

Additional file 7
A. rpoB marker
s
A. 16S marker

### Rarefaction curves obtained by Illumina-amplicon sequencing of rpoB (A) and 16S (B) markers in 15 mock community samples

#### Slide 1
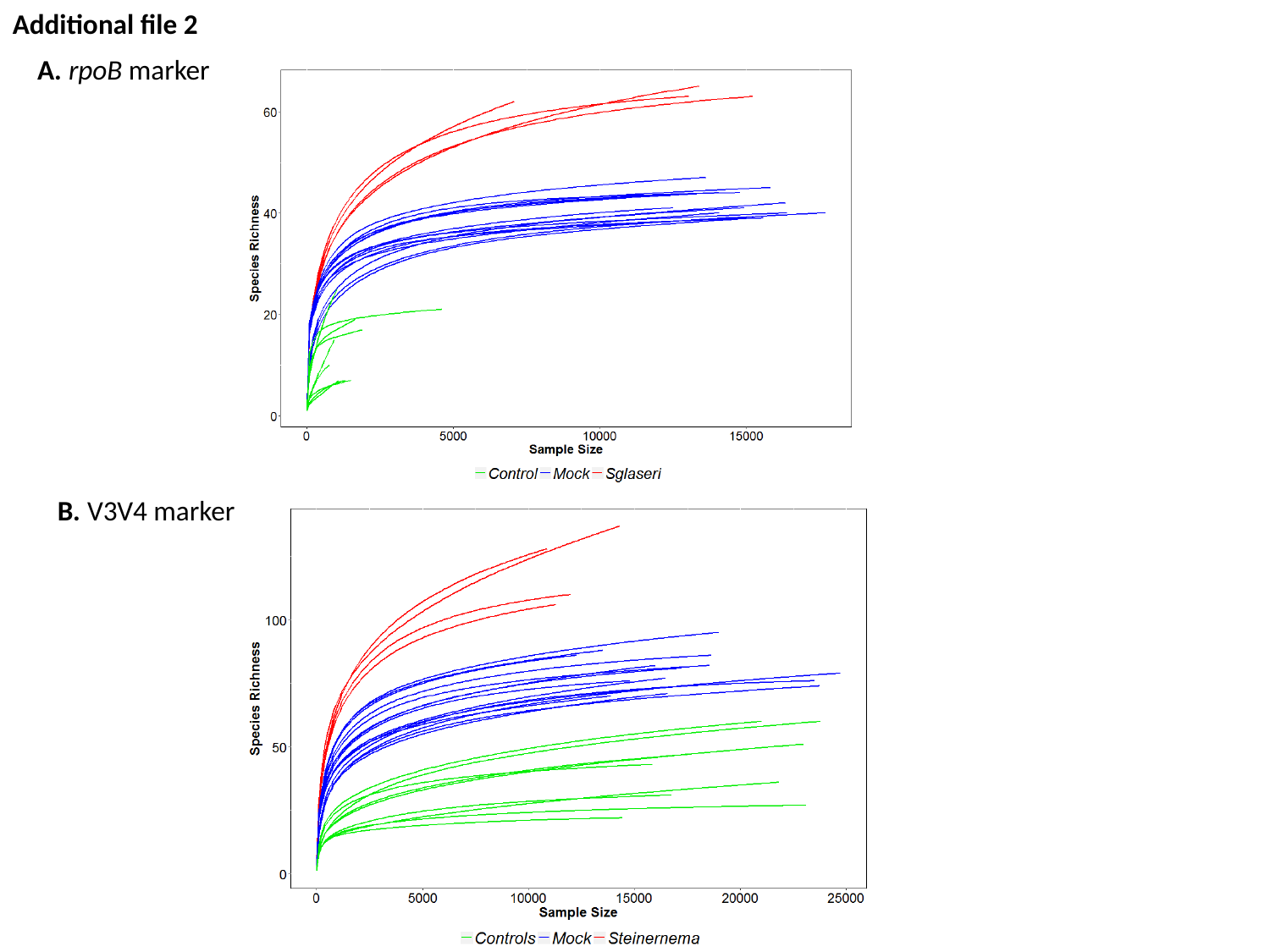

Additional file 2
A. rpoB marker
B. V3V4 marker
