## Supplementary material for "*rpoB*, a promising marker for analyzing the diversity of bacterial communities by amplicon sequencing": Comparison of the expected bacterial composition and the observed OTU composition generated by Illumina-amplicon rpoB sequencing for the mock1

### Slide 1
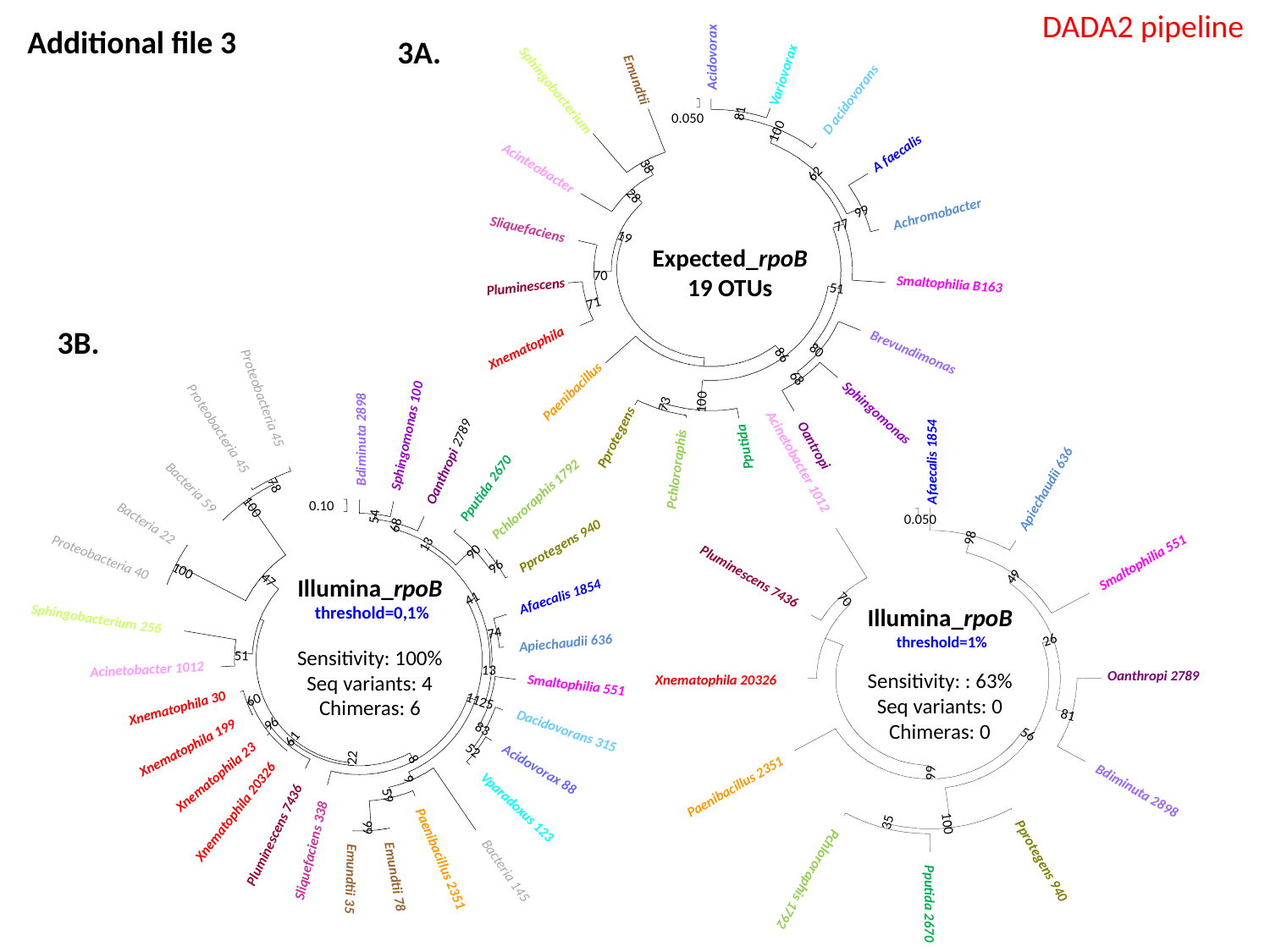

DADA2 pipeline
Acidovorax
Variovorax
Emundtii
Sphingobacterium
D acidovorans
81
0.050
100
A faecalis
38
Acinteobacter
62
28
99
Achromobacter
77
Sliquefaciens
19
70
Smaltophilia B163
Pluminescens
51
71
Xnematophila
80
Brevundimonas
86
68
Paenibacillus
100
73
Sphingomonas
Pprotegens
Oantropi
Pputida
Pchlororaphis
3A.
Expected_rpoB
19 OTUs
Additional file 3
3B.
Proteobacteria 45
Proteobacteria 45
Sphingomonas 100
Bdiminuta 2898
Oanthropi 2789
78
Bacteria 59
Pputida 2670
Pchlororaphis 1792
0.10
100
54
Bacteria 22
68
13
Pprotegens 940
90
Proteobacteria 40
96
100
47
Afaecalis 1854
41
Sphingobacterium 256
74
Apiechaudii 636
51
Acinetobacter 1012
13
Smaltophilia 551
11
60
25
Xnematophila 30
96
83
Dacidovorans 315
61
Xnematophila 199
52
22
8
Acidovorax 88
Xnematophila 23
9
59
Vparadoxus 123
Xnematophila 20326
99
Pluminescens 7436
Sliquefaciens 338
Paenibacillus 2351
Bacteria 145
Emundtii 78
Emundtii 35
Acinetobacter 1012
Afaecalis 1854
Apiechaudii 636
0.050
98
Smaltophilia 551
Pluminescens 7436
49
70
26
Oanthropi 2789
Xnematophila 20326
81
56
99
Paenibacillus 2351
Bdiminuta 2898
35
100
Pprotegens 940
Pchlororaphis 1792
Pputida 2670
Illumina_rpoB
 threshold=0,1%
Sensitivity: 100%
Seq variants: 4
Chimeras: 6
Illumina_rpoB
 threshold=1%
Sensitivity: : 63%
Seq variants: 0
Chimeras: 0
