## Supplementary material for "*rpoB*, a promising marker for analyzing the diversity of bacterial communities by amplicon sequencing": Comparison of expected bacterial composition and the observed OTU composition obtained by Illumina-amplicon 16S rRNA gene sequencing for the mock1

### Slide 1
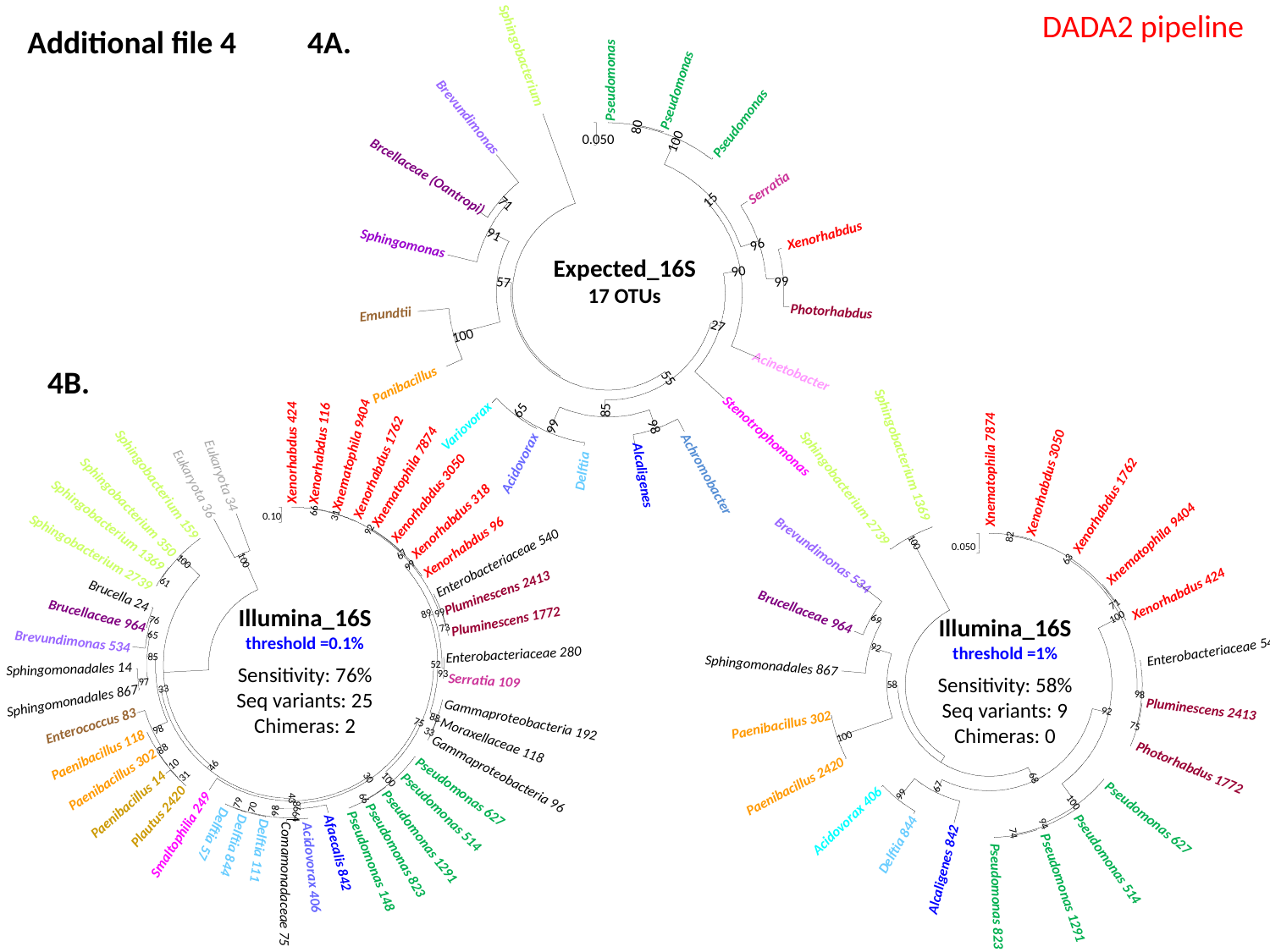

DADA2 pipeline
Sphingobacterium
Pseudomonas
Pseudomonas
Brevundimonas
Pseudomonas
80
0.050
100
Brcellaceae (Oantropi)
Serratia
15
71
91
Xenorhabdus
Sphingomonas
96
90
99
57
Photorhabdus
Emundtii
27
100
Acinetobacter
55
Panibacillus
65
85
Variovorax
99
Acidovorax
Delftia
98
Stenotrophomonas
Achromobacter
Alcaligenes
Additional file 4
4A.
Expected_16S
17 OTUs
4B.
Sphingobacterium 1369
Xnematophila 7874
Xenorhabdus 3050
Sphingobacterium 2739
Xenorhabdus 1762
82
Xnematophila 9404
100
0.050
Brevundimonas 534
63
Xenorhabdus 424
71
Brucellaceae 964
100
69
92
Enterobacteriaceae 540
Sphingomonadales 867
58
98
Pluminescens 2413
92
Paenibacillus 302
75
100
Photorhabdus 1772
68
Paenibacillus 2420
67
99
100
Pseudomonas 627
Acidovorax 406
94
74
Delftia 844
Pseudomonas 514
Alcaligenes 842
Pseudomonas 1291
Pseudomonas 823
Xenorhabdus 424
Xenorhabdus 116
Xnematophila 9404
Xenorhabdus 1762
Eukaryota 34
Xnematophila 7874
Eukaryota 36
Sphingobacterium 159
Xenorhabdus 3050
Sphingobacterium 350
66
31
0.10
Xenorhabdus 318
Sphingobacterium 1369
92
Xenorhabdus 96
Sphingobacterium 2739
67
Enterobacteriaceae 540
100
100
99
61
Pluminescens 2413
Brucella 24
99
89
Brucellaceae 964
Pluminescens 1772
76
73
65
Brevundimonas 534
Enterobacteriaceae 280
85
52
Sphingomonadales 14
93
Serratia 109
97
33
Sphingomonadales 867
88
Gammaproteobacteria 192
75
Enterococcus 83
98
33
Moraxellaceae 118
88
Paenibacillus 118
10
46
Gammaproteobacteria 96
31
Paenibacillus 302
30
100
Pseudomonas 627
43
66
Paenibacillus 14
79
86
70
Pseudomonas 514
98
Plautus 2420
64
Smaltophilia 249
Delftia 57
Pseudomonas 1291
Delftia 844
Pseudomonas 823
Delftia 111
Afaecalis 842
Pseudomonas 148
Acidovorax 406
Comamonadaceae 75
Illumina_16S
threshold =0.1%
Sensitivity: 76%
Seq variants: 25
Chimeras: 2
Illumina_16S
threshold =1%
Sensitivity: 58%
Seq variants: 9
Chimeras: 0
