## Supplementary material for "*rpoB*, a promising marker for analyzing the diversity of bacterial communities by amplicon sequencing": Comparison of the observed and expected relative abundances of the bacterial communities obtained by Illumina-amplicon rpoB(A) and 16S(B) sequencing

### Slide 1
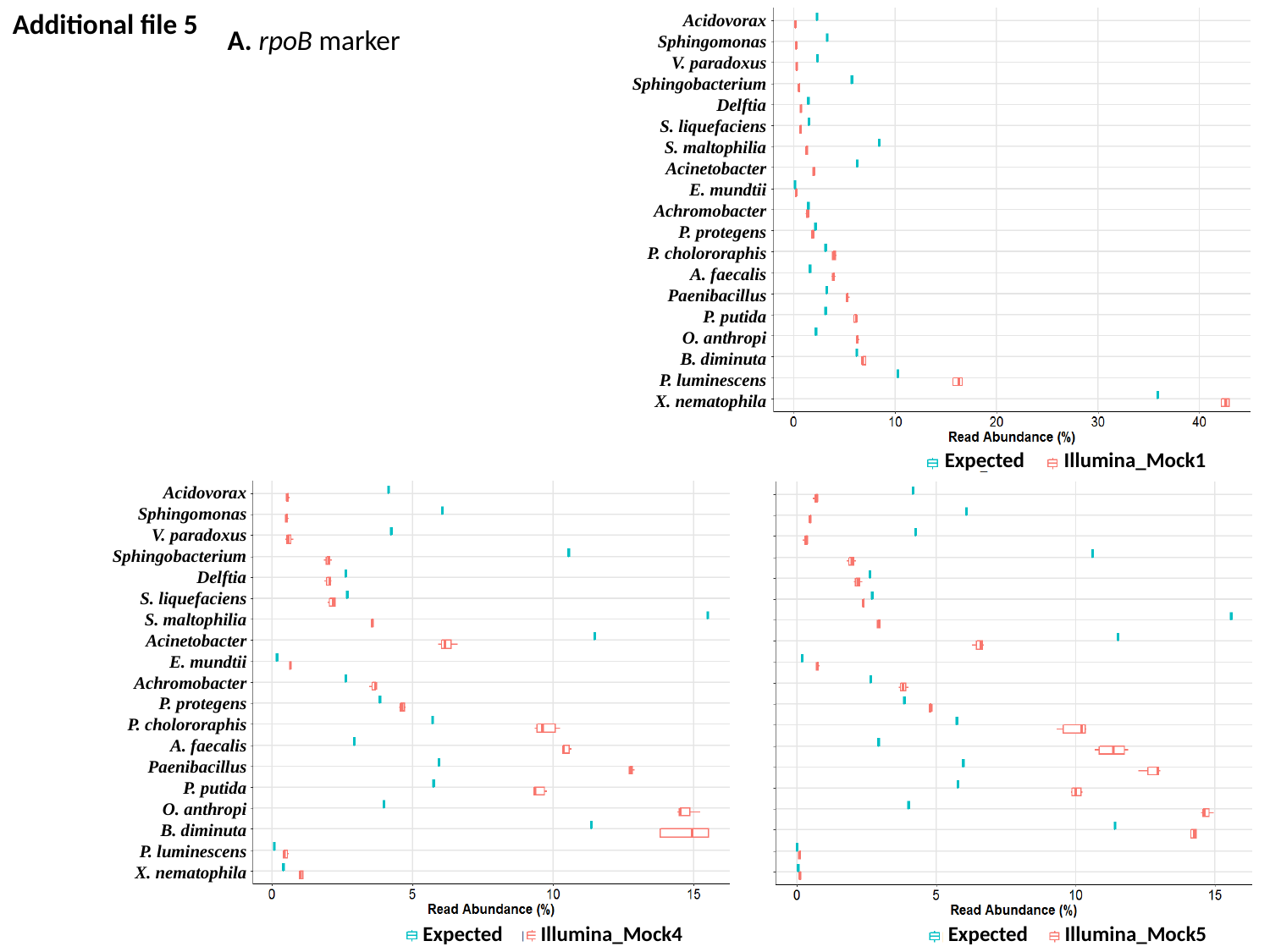

Additional file 5
Acidovorax
Sphingomonas
V. paradoxus
Sphingobacterium
Delftia
S. liquefaciens
S. maltophilia
Acinetobacter
E. mundtii
Achromobacter
P. protegens
P. cholororaphis
A. faecalis
Paenibacillus
P. putida
O. anthropi
B. diminuta
P. luminescens
X. nematophila
Expected
Illumina_Mock1
A. rpoB marker
Acidovorax
Sphingomonas
V. paradoxus
Sphingobacterium
Delftia
S. liquefaciens
S. maltophilia
Acinetobacter
E. mundtii
Achromobacter
P. protegens
P. cholororaphis
A. faecalis
Paenibacillus
P. putida
O. anthropi
B. diminuta
P. luminescens
X. nematophila
Expected
Illumina_Mock4
Expected
Illumina_Mock5

### Slide 2
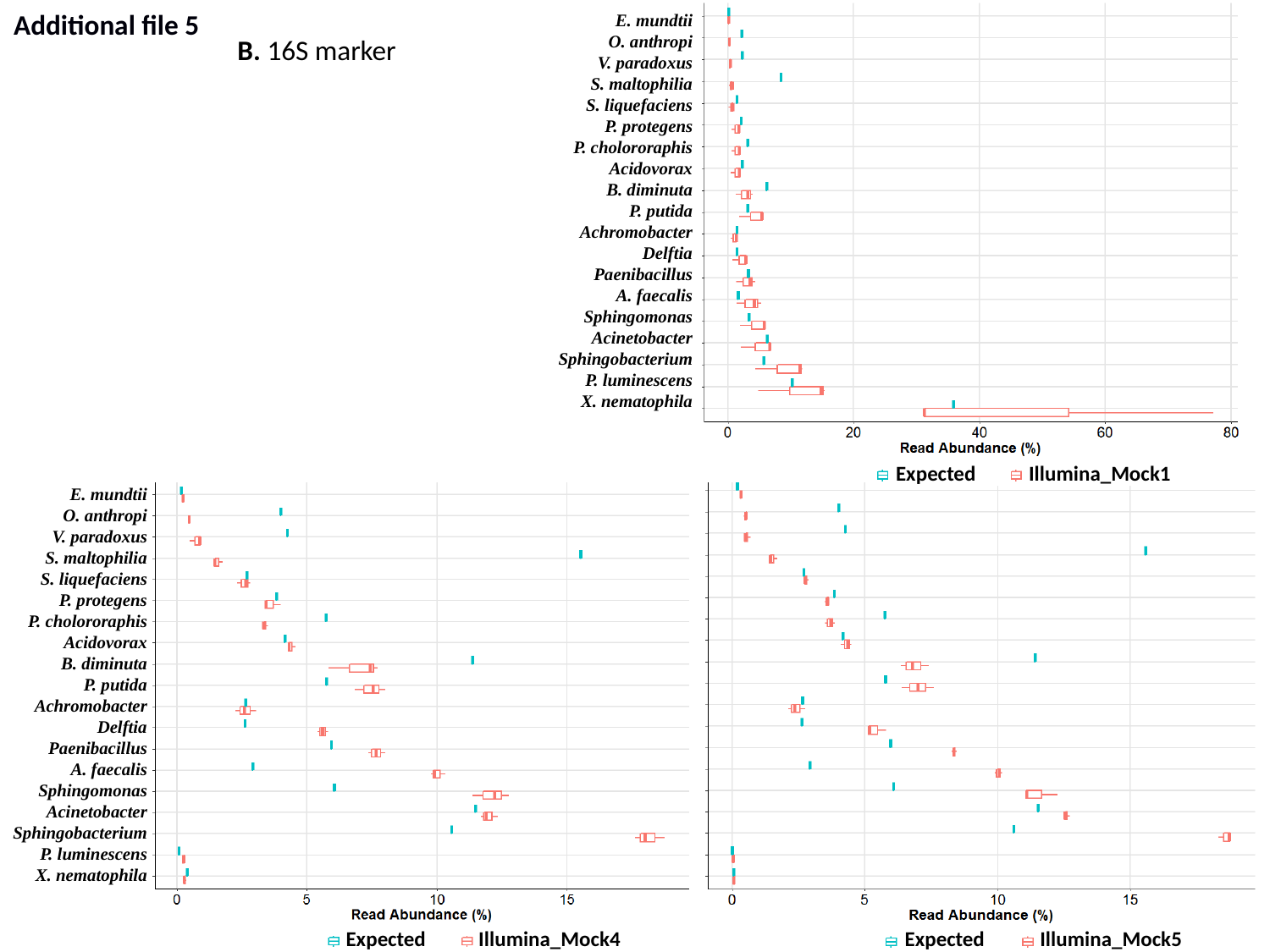

Additional file 5
E. mundtii
O. anthropi
V. paradoxus
S. maltophilia
S. liquefaciens
P. protegens
P. cholororaphis
Acidovorax
B. diminuta
P. putida
Achromobacter
Delftia
Paenibacillus
A. faecalis
Sphingomonas
Acinetobacter
Sphingobacterium
P. luminescens
X. nematophila
B. 16S marker
Expected
Illumina_Mock1
E. mundtii
O. anthropi
V. paradoxus
S. maltophilia
S. liquefaciens
P. protegens
P. cholororaphis
Acidovorax
B. diminuta
P. putida
Achromobacter
Delftia
Paenibacillus
A. faecalis
Sphingomonas
Acinetobacter
Sphingobacterium
P. luminescens
X. nematophila
Expected
Illumina_Mock4
Expected
Illumina_Mock5
