## Supplementary material for "*rpoB*, a promising marker for analyzing the diversity of bacterial communities by amplicon sequencing": Sequence reads and OTU numbers obtained by Illumina-amplicon sequencing of rpoB and 16S markers for the nematode samples (four replicates).

**Additional file 6**

| Sample ID | Reads after filtering | | | | Number of OTUs (reads ≥ 0.1%) | |
| --- | --- | --- | --- | --- | --- | --- |
|  | FROGS | | DADA2 | | FROGS | DADA2 |
| rpoB-SK39-rep1 | 7,062 | | 8134 | | 30 | 31 |
| rpoB-SK39-rep2 | 13,036 | | 15240 | | 34 | 36 |
| rpoB-SK39-rep3 | 13,402 | | 15468 | | 31 | 33 |
| rpoB-SK39-rep4 | 15,219 | | 17857 | | 30 | 31 |
| 16S-SK39-rep1 | 10,863 | | 9794 | | 53 | 52 |
| 16S-SK39-rep2 | 13,141 | | 11257 | | 58 | 51 |
| 16S-SK39-rep3 | 11,294 | | 10023 | | 50 | 48 |
| 16S-SK39-rep4 | 12,000 | | 10581 | | 55 | 47 |
