## Supplementary material for "*rpoB*, a promising marker for analyzing the diversity of bacterial communities by amplicon sequencing": Comparison of the bacterial compositions obtained by Illumina sequencing of rpoB and 16S rRNA for the nematode S. glaseri SK39 (DADA2 process).

### Slide 1
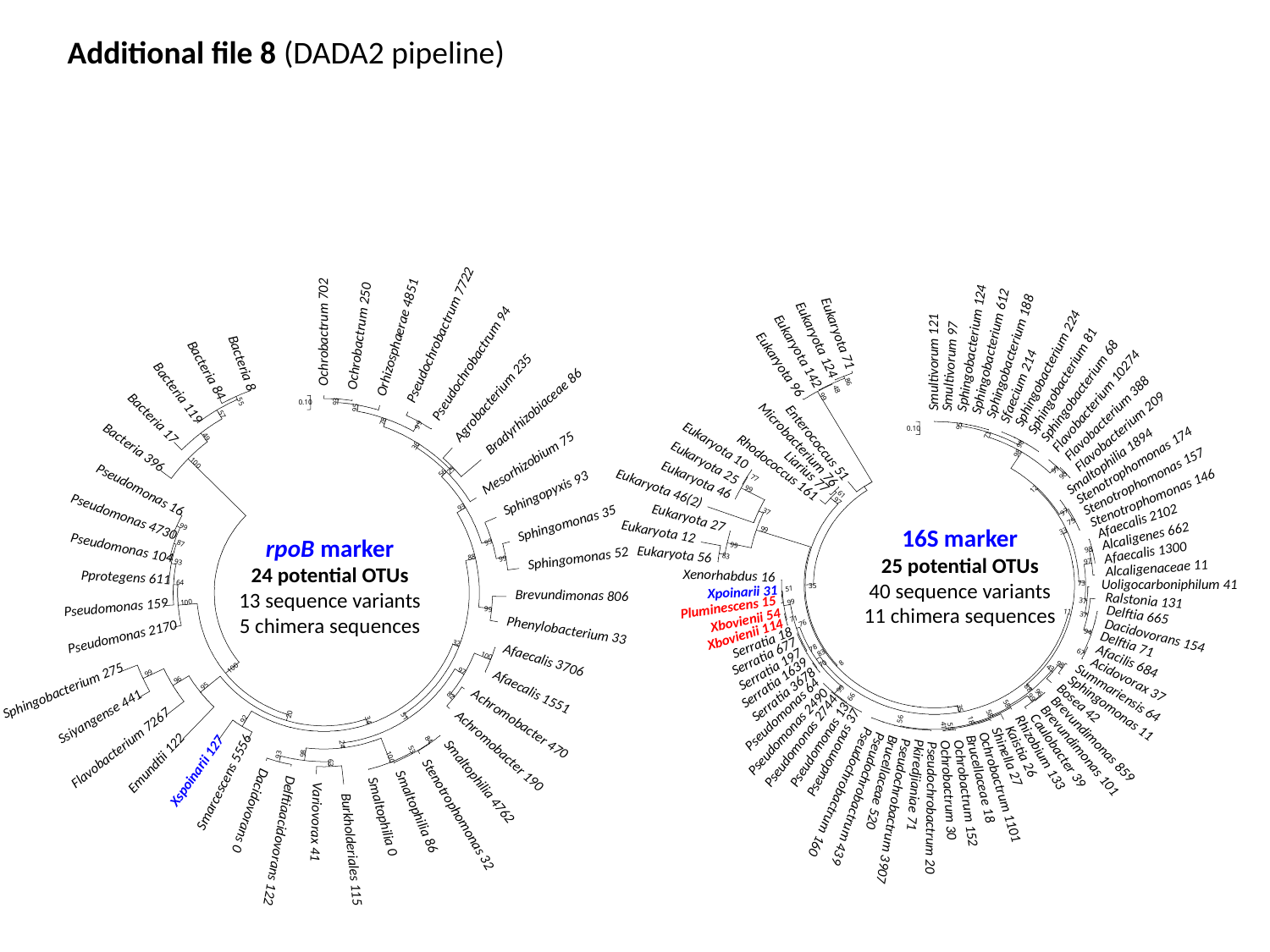

Additional file 8 (DADA2 pipeline)
Ochrobactrum 702
Pseudochrobactrum 7722
Ochrobactrum 250
Orhizosphaerae 4851
Pseudochrobactrum 94
Agrobacterium 235
93
Bradyrhizobiaceae 86
95
78
99
36
Mesorhizobium 75
25
56
Pseudomonas 16
Sphingopyxis 93
93
Pseudomonas 4730
Sphingomonas 35
99
99
Pseudomonas 104
Sphingomonas 52
88
99
Pprotegens 611
Brevundimonas 806
Pseudomonas 159
99
Phenylobacterium 33
Pseudomonas 2170
32
Afaecalis 3706
100
97
99
96
Sphingobacterium 275
95
Afaecalis 1551
82
Ssiyangense 441
20
51
92
Achromobacter 470
34
84
Flavobacterium 7267
24
Achromobacter 190
53
98
93
100
Emundtii 122
63
Xspoinarii 127
Smarcescens 5556
Smaltophilia 4762
Dacidovorans 0
Smaltophilia 86
Stenotrophomonas 32
Smaltophilia 0
Variovorax 41
Delftiaacidovorans 122
Burkholderiales 115
Bacteria 8
Bacteria 84
Bacteria 119
55
0.10
Bacteria 17
57
49
Bacteria 396
100
87
93
64
100
100
Sphingobacterium 124
Sphingobacterium 612
Sphingobacterium 188
Smultivorum 121
Smultivorum 97
Sphingobacterium 224
Sphingobacterium 81
Sfaecium 214
Sphingobacterium 68
Flavobacterium 10274
Flavobacterium 388
95
Flavobacterium 209
71
98
98
Smaltophilia 1894
Stenotrophomonas 174
99
90
Stenotrophomonas 157
12
Stenotrophomonas 146
97
Afaecalis 2102
75
33
Alcaligenes 662
Afaecalis 1300
98
97
Alcaligenaceae 11
Uoligocarboniphilum 41
73
Ralstonia 131
37
Delftia 665
37
Dacidovorans 154
94
Delftia 71
67
Afacilis 684
98
43
Acidovorax 37
Summariensis 64
96
99
Bosea 42
Sphingomonas 11
58
58
56
18
41
55
Brevundimonas 859
Brevundimonas 101
Caulobacter 39
Kaistia 26
Rhizobium 133
Shinella 27
Brucellaceae 18
Brucellaceae 520
Pkiredjianiae 71
Ochrobactrum 1101
Ochrobactrum 30
Pseudochrobactrum 160
Ochrobactrum 152
Pseudochrobactrum 439
Pseudochrobactrum 20
Pseudochrobactrum 3907
Eukaryota 71
Eukaryota 124
Eukaryota 142
Eukaryota 96
86
48
99
0.10
Enterococcus 51
Microbacterium 76
Eukaryota 10
Eukaryota 25
Rhodococcus 161
Liarius 77
Eukaryota 46
77
Eukaryota 46(2)
99
61
97
37
Eukaryota 27
Eukaryota 12
99
99
Eukaryota 56
83
Xenorhabdus 16
35
Xpoinarii 31
51
99
Pluminescens 15
11
Xbovienii 54
71
76
Xbovienii 114
Serratia 18
78
Serratia 677
83
8
70
Serratia 197
Serratia 1639
88
99
Serratia 3678
66
35
Pseudomonas 64
Pseudomonas 2490
Pseudomonas 2744
Pseudomonas 13
Pseudomonas 37
16S marker
25 potential OTUs
40 sequence variants
11 chimera sequences
rpoB marker
24 potential OTUs
13 sequence variants
5 chimera sequences
