## Supplementary material for "*rpoB*, a promising marker for analyzing the diversity of bacterial communities by amplicon sequencing": List of biological materials used in the experimental study

**Additional file 9**

**A. List of the bacterial strain isolates used to generate the mock communities**

| **Species** | **Strain** | **Source of isolation** | **Reference** |
| --- | --- | --- | --- |
| *Xenorhabdus nematophila* | F1 | Symbiont of *S. carpocapsae* SK27 (hanging drop isolation technique) | [1, 2] |
| *Photorhabdus luminescens* | TT01 | Symbiont of *H. bacteriophora TT01(hanging drop isolation technique)* | [3, 4] |
| *Pseudomonas protegens* | SW4 | *G. mellonella cadaver after IJ infestation with S. weiseri* 583 | This study |
| *Pseudomonas chlororaphis* | SK39 ApoA | *G. mellonella* cadaver after IJ infestation with *S. glaseri* SK39 | This study |
| *Pseudomonas putida* | SW5 | *G. mellonella* cadaver after IJ infestation with *S. weiseri* 583 | This study |
| *Stenotrophomonas maltophilia* | B163 | *G. mellonella* cadaver after IJ infestation with *S. carpocapsae SK27* | This study |
| *Achromobacter sp.* | SK39 ApoC | *G. mellonella* cadaver after IJ infestation with *S. glaseri* SK39 | This study |
| *Alcaligenes faecalis* | SC | Isolated after the crushing of *S. carpocapsae* SK27 IJs | This study |
| *Ochrobactrum anthropi* | SW2 | *G. mellonella* cadaver after IJ infestation with *S. weiseri* 583 | This study |
| *Acinetobacter* | FR211 A | *G. mellonella* cadaver after IJ infestation with *S. affine* | This study |
| *Serratia liquefaciens* | TUR03-2-2 | Isolated from *S. weiserii* 583 (hanging drop isolation technique) | This study |
| *Delftia sp.* | MW8A-4 | Isolated from storage medium of a batch of *S. tanzaniensis* IJs | This study |
| *Variovorax sp.* | BS 3-2-1 | Isolated after the crushing of *H. bacteriophora* BS 3.2 IJs | This study |
| *Acidovorax sp.* | 173 | Isolated after crushing of *S. xueshanense* Dequen IJs | This study |
| *Sphingomonas sp.* | BG | Isolated after the crushing of *H. bacteriophora* CS3-2 IJs | This study |
| *Brevundimonas sp.* | TCH07-Apo2-4 | Isolated from *S. weiserii 583* by the hanging drop technique | This study |
| *Sphingobacterium sp.* | FR211C | *G. mellonella* cadaver after IJ infestation with *S. affine* | This study |
| *Paenibacillus sp.* | SK72-2 | *G. mellonella* cadaver after IJ infestation with *S. glaseri* SK72 | This study |
| *Enterococcus mundtii* | SG4 | *G. mellonella* cadaver after IJ infestation with *S. glaseri* SK39 | This study |

**B. List of the primers used in this study**

| **Primer label** | **Sequence (5’-3’)** | **Utilization** | **Reference** |
| --- | --- | --- | --- |
| 16S_F | GAAGAGTTTGATCATGGCTC | 16S rRNA gene Sanger sequencing | [5] |
| 16S_R | AAGGAGGTGATCCAGCCGCA |  |  |
| Univ_rpoB_F_deg | **GGYTWYGAAGTNCGHGACGTDCA** | *rpoB* gene fragment Sanger sequencing | This study |
| Univ_rpoB_R_deg | **TGACGYTGCATGTTBGMRCCCATMA** |  |  |
| F343 | CTTTCCCTACACGACGCTCTTCCGATCTTA**CGGRAGGCAGCAG** | V3V4 Illumina-amplicon sequencing | [6] |
| R784 | GGAGTTCAGACGTGTGCTCTTCCGATCTTA**CCAGGGTATCTAATCCT** |  |  |
| Univ_rpoB_F_deg | CTTTCCCTACACGACGCTCTTCCGATCT**GGYTWYGAAGTNCGHGACGTDCA** | *rpoB* Illumina-amplicon sequencing | This study |
| Univ_rpoB_R_deg | GGAGTTCAGACGTGTGCTCTTCCGATCT**TGACGYTGCATGTTBGMRCCCATMA** |  |  |
